# Modified N-linked glycosylation status predicts trafficking defective human Piezo1 channel mutations

**DOI:** 10.1101/2020.11.30.404962

**Authors:** Jinyuan Vero Li, Chai-Ann Ng, Delfine Cheng, Mingxi Yao, Yang Guo, Ze-Yan Yu, Yogambha Ramaswamy, Lining Arnold Ju, Philip W Kuchel, Michael P Feneley, Diane Fatkin, Charles D Cox

## Abstract

Mechanosensitive channels are integral membrane proteins that sense mechanical stimuli. Like all membrane proteins, they pass through biosynthetic quality control in the endoplasmic reticulum and Golgi that results in them reaching their destination at the plasma membrane. Here we show that N-linked glycosylation of two highly conserved asparagine residues in the ‘cap’ region of mechanosensitive Piezo1 channels are necessary for the mature protein to reach the plasma membrane. Both mutation of these asparagines (N2294Q/N2331Q) and treatment with an enzyme that hydrolyses N-linked oligosaccharides (PNGaseF) eliminates the fully glycosylated mature Piezo1 protein. The N-glycans in the cap are a pre-requisite for higher-order glycosylation in the ‘propeller’ regions, which are present in loops that are essential for mechanotransduction. Importantly, trafficking-defective Piezo1 variants linked to generalized lymphatic dysplasia and bicuspid aortic valve display reduced fully N-glycosylated protein. The higher order glycosylation status *in vitro* correlates with efficient membrane trafficking and will aid in determining the functional impact of Piezo1 variants of unknown significance.

## Introduction

The Piezo family of ion channels has only two members, Piezo1 and Piezo2^1,2^. These are large membrane proteins with more than 30 transmembrane helices that decode mechanical cues. Piezo1 in particular appears to be a central mechanotransducer in the cardiovascular system^3,4^ and is sensitive to membrane forces^2,5–7^ and thus lipid composition^8–12^. Like all other integral membrane proteins, Piezo1 channels undergo biosynthetic quality control in the endoplasmic reticulum (ER) and Golgi^13^. This study focussed on exactly how these processes regulate the membrane expression of Piezo1.

All membrane proteins undergo folding and maturation in the ER and Golgi, where N-linked glycosylation usually fulfils a critical role in biosynthetic quality control^13,14^. N-linked glycosylation is the process by which oligosaccharides are covalently attached to asparagine residues in proteins. This process begins with the co-translational addition of core-glycans in the ER and culminates in the processing and addition of higher order glycans in the Golgi prior to vesicular transit to the plasma membrane. The higher order glycans can be high mannose, complex, or hybrid glycans, depending on which types of carbohydrates are added^13,15^. Once the glycans are added to the membrane protein, they alter their folding and stability^16,17^. In many cases, the glycans act as a ‘quality control stamp’ certifying the folding status of the membrane protein during its biosynthesis.

It is particularly evident that N-linked glycosylation is important in the biosynthetic quality control of plasma membrane ion channels. The hERG K^+^ channel (K_v11.1_)^18,19^, cystic fibrosis transmembrane conductance regulator (CFTR)^20^ and polycystic kidney disease proteins^21^ have all been studied in this regard. Variants in these ion channels cause channelopathies that arise from two broad mechanisms: 1) a functional defect in the channel; or 2) aberrant plasma membrane trafficking. In fact, most disease-causing variants are aberrantly trafficked. For example, in the case of K_v11.1_ mutations, ~80% of variants linked to long QT syndrome type II (LQTS2) ameliorate trafficking^22^. The N-linked glycosylation of K_v11.1_ produces a characteristic pattern on a Western blot consisting of a fully glycosylated protein of larger molecular size (~155 kDa) and a core glycosylated protein that is smaller (~135 kDa)^18,19,22–25^. The upper band (larger molecular size) is a surrogate of the mature membrane protein^17^ and as a result, has been used extensively to interrogate variants of unknown significance in K_v11.1_^22,24,26^. Thus, trafficking-defective mutants lack the mature form, and rather than showing a double band appearance on a Western blot they have a single band that represents the core-glycosylated protein. Given that variants of Piezo1 have been linked to disease (e.g., generalized lymphatic dysplasia^27,28^ and bicuspid aortic valve^29^) we explored whether human Piezo1 undergoes N-linked glycosylation and whether this serves as a predictor of efficient membrane trafficking.

Here we report on the molecular events that occur during the trafficking of Piezo1 channels and we showed that N-linked glycosylation was critical in the biosynthetic quality control of this channel. In particular, we demonstrated a novel mechanism whereby two highly conserved asparagines in the cap (N2294Q and N2331Q), which are essential for normal trafficking of Piezo1 to the plasma membrane, dictated the glycosylation status of the propeller domains that extend out from the central ion conducting pore^30–32^. Also, higher order glycosylation of the N-terminus was seen to be a surrogate for the mature membrane protein during the trafficking of various human Piezo1 variants. These findings will be useful when assessing the effects of mutations generated during structure-function studies, or when exploring disease-linked variants. As a proof of concept, we showed that loss-of-function Piezo1 variants such as G2029R that are known to be trafficking-defective^28^ lack higher order N-glycosylation when expressed *in vitro*. Furthermore, we revealed that some loss-of-function variants that affect trafficking display a dominant negative effect on function and retard the trafficking of the wild-type protein. These findings have broad significance for future studies of diseases resulting from *PIEZO1* variants.

## Methods

### Cell lines

Piezo1^-/-^ HEK293T cells^28^ were a gift from Dr Ardem Patapoutian (The Scripps Research Institute, La Jolla, CA, USA); HEK293 (R78007, Scientific, Waltham, MA, USA) was a gift from Dr Nicola Smith (UNSW, Sydney, NSW, Australia); HEK293S GnT1^-/-^ cells were a gift from Dr Jamie Vandenberg (VCCRI, Sydney, NSW, Australia); and human BJ-5ta-hTERT foreskin fibroblasts were provided by Dr Michael Sheetz (UTMB, Galveston, TX, USA). Cell lines were not authenticated and were not listed in the database of commonly misidentified cell lines maintained by ICLAC (http://iclac.org) and NCBI Biosample (http://www.ncbi.nlm.nih.gov/biosample). All cell lines were confirmed to be mycoplasma free.

### Mutagenesis

Site directed mutagenesis of human and mouse Piezo1 was undertaken using a custom protocol with the high-fidelity polymerase PfuUltra. The mouse C-terminal GFP fusion construct was generated by deleting the IRES sequence using site-directed mutagenesis from the pcDNA3.1 IRES GFP construct of mouse Piezo1 (Provided by Dr Ardem Patapoutian). Primers for point mutations and fusion protein generation are listed in Table1.

**Table 1.**
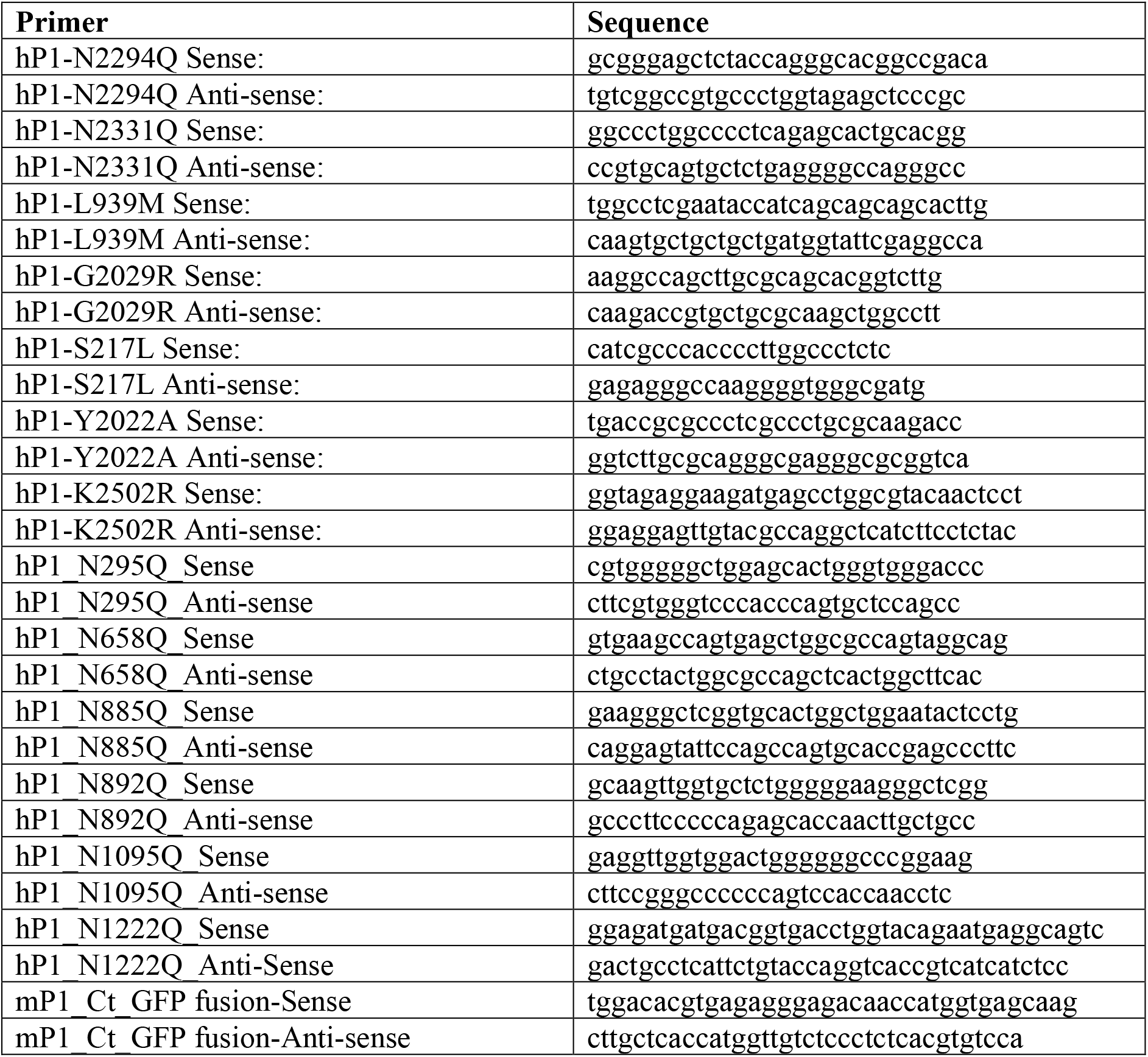
List of primers for site-directed mutagenesis of human (hP1) and mouse (mP1) Piezo1.

### Western blotting

Cells were cultured in Dulbecco’s modified Eagle medium (DMEM; Sigma-Aldrich, St. Louis, MO, USA) supplemented with 10% v/v foetal bovine serum (ThermoFisher Scientific, Waltham, MA, USA) and incubated at 37°C with 5% CO_2_. GFP or mCherry-fused WT and mutant human Piezo1 cloned from HEK cells^33^, or GFP-fused mouse Piezo1 were transfected into HEK293T Piezo1^-/-^ cells, HEK293S GnT1^-/-^ or HEK293 cells (ThermoFisher Scientific, Cat. No. R78007) using Lipofectamine 3000 transfection reagent (ThermoFisher Scientific) with 800 ng of DNA (400 ng WT and mutants, respectively, for co-expression transfection). The medium was changed 24 h after transfection. Cells were harvested 72 h after transfection and solubilized in radio-immunoprecipitation assay buffer (RIPA) buffer [Tris buffer 10 mM, ethylenediaminetetraacetic acid (EDTA) 1 mM, NaCl 140 mM, in (% w/v): Sodium deoxycholate 0.1, SDS 0.1, Triton X-100 1.0, pH 7.2] supplemented with 1 × EDTA-free protease inhibitor cocktail tablets (Sigma-Aldrich), 1 mM (phenylmethylsulfonyl fluoride) PMSF, 2 mM tris(2-carboxyethyl)phosphine (TCEP), and 1 mM *N*-ethylmaleimide (NEM) for 10 min on a rotating wheel at 4°C. Cell lysates were cleared by centrifugation at 13,000 × g at 4°C for 20 min. Transfection efficiency was estimated, and the relative Piezo1 concentration was determined from the intensity of fluorescence of the lysate using a PHERAstarFS microplate reader (BMG LABTECH, Ortenberg, Germany).

For PNGaseF (peptide N-glycosidase F, EC 3.5.1.52) digestion, 5% v/v PNGaseF (New England Biolabs, Ipswich, MA, USA) and 50 mM NEM were added to lysates mixed with 1 × GlycoBuffer and incubated on ice for 1 h. Lysates that were undigested or digested with PNGaseF were then mixed with SDS-PAGE sample buffer, and loaded and run on a 3-8% Tris-Acetate gel (Thermo Fisher Scientific) before being transferred to a nitrocellulose membrane (Bio-Rad, Hercules, CA, USA).

For quantitative Western blot analysis, GFP-fused Piezo1 was probed with a rabbit monoclonal anti-GFP antibody (Santa Cruz Biotechnology, Dallas, TX, USA; 1:5,000 dilution); mCherry-fused Piezo1 protein was probed with rat monoclonal anti-mCherry antibody (Clone 16D7, ThermoFisher Scientific; 1:1,000 dilution); the native human Piezo1 channel was probed using a mouse monoclonal anti-Piezo1 antibody (Cat# NBP2-75617, Novus Biologicals, Centennial, CO, USA; 1:1,000 dilution); mouse anti-α-actinin antibody (Santa Cruz Biotechnology; 1:5,000 dilution) or anti-α-tubulin (Clone DM1A, Sigma Aldrich, T9026) antibody was added simultaneously for a loading comparison followed by anti-rabbit IRDye680 at 1:20,000, anti-rat

IRDye8OO at 1:10,000 and anti-mouse IRDye800 at 1:20,000 (Li-Cor) to enable quantification with the LI-COR Odyssey system (LI-COR Biotechnology, Lincoln, NE, USA). Image studio (LI-COR Biotechnology) was used to calculate the ratio of the fully glycosylated upper band (FG)/core-glycosylated lower band (CG) in a manner that was similar to previous reports^26^.

### Electrophysiology

Transiently transfected Piezo1^-/-^ HEK293T cells were plated on 35 mm dishes for patch clamp analysis. The extracellular solution for cell-attached patches contained high K^+^ to zero the membrane potential; it consisted of 90 mM potassium aspartate, 50 mM KCl, 1 mM MgCl_2_ and 10 mM HEPES (pH 7.2) adjusted with 5 M KOH. The pipette solution contained either 140 mM CsCl or 140 mM NaCl with 10 mM HEPES (pH 7.2) adjusted with the respective hydroxide. Ethylene glycol-bis(β-aminoethyl etlier)-*N,N,N′,N′*-tetraacetic acid (EGTA) was added to control levels of free pipette (extracellular) Ca^2+^ using the online EGTA calculator—Ca-EGTA Calculator TS v1.3—Maxchelator. Negative pressure was applied to patch pipettes using a High Speed Pressure Clamp-1 (ALA Scientific Instruments, Farmingdale, NY, USA) and recorded in millimetres of mercury (mmHg) using a piezoelectric pressure transducer (WPI, Sarasota, FL, USA). Borosilicate glass pipettes (Sigma-Aldrich) were pulled with a vertical pipette puller (PP-83, Narashige, Tokyo, Japan) to produce electrodes with a resistance of 1.8-2.2MΩ. Single-channel Piezo1 currents were amplified using an AxoPatch 200B amplifier (Axon Instruments, Union City, CA, USA), and data were sampled at a rate of 10 kHz with 1 kHz filtration, and analysed using pCLAMP10 software (Axon Instruments). The Boltzmann distribution function was used to describe the dependence of mesoscopic Piezo1 channel currents and open probability, respectively, on the negative pressure applied to patch pipettes. Boltzmann plots were obtained by fitting open probability *P*_o_~*I/I*_max_ versus negative pressure using *P*_o_/(1−*P*_o_) = exp [*α* (*P*-*P*_1/2_)], where *P* is the negative pressure (suction) in mm Hg, *P*_1/2_ is the negative pressure at which *P*o = 0.5, and *α* (mm Hg)^-1^ is the slope of the plot of ln [*P*_o_/(1−*P*_o_)] versus (*P*−*P*_1/2_), reflecting the channels’ mechanosensitivity.

Single-channel amplitudes were measured by Gaussian fits (Clampfit; Axon Instruments) on an all-points-histogram of current amplitudes over a 2 s period exhibiting only single-channel openings. Conductance was then calculated by regressing a line on the graph of current amplitude vs holding potential at five separate voltages.

### Immunogold labelling and electron microscopy

Cells were grown to 70-80% confluency on fibronectin-coated coverslips before being fixed with 4% w/v paraformaldehyde (PFA) in 100 mM Sorensen’s phosphate buffer (pH 7.2) for 20 min. Piezo1 localisation was detected using electron microscopy (EM) with immunogold labelling using nanogold followed by silver enhancement. The protocol was adapted from Biazik et al^34^ as follows: Free aldehyde was quenched with 0.1 M glycine for 20 min and the cell membranes were permeabilized with 0.005% w/v saponin (containing 0.1% w/v bovine serum albumin in 1 × PBS) for 8 min. The samples were then incubated with mouse monoclonal anti-Piezo1 antibody (1:60 dilution, Cat# NBP2-75617, Novus Biologicals) overnight at 4°C. The next day, the samples were washed and incubated with a secondary antibody that was conjugated with 1.4 nm nanogold (1:60 dilution, Cat# 2002-0.5 mL, Nanoprobes, Yaphank, NY, USA) for 1 h. The labelled cells were then fixed in 2.5% w/v glutaraldehyde for 10 min and quenched with 0.1 M glycine for 20 min. The nanogold was silver enhanced for 7 min using an HQ silver enhancement kit (Cat# 2012-45 mL, Nanoprobes). The silver was further stabilised by gold toning that involved 15 min incubation in 2% w/v sodium acetate, 10 min incubation in 0.05% w/v gold (III) chloride trihydrate (on ice) and 10 min incubation in 0.3% w/v sodium thiosulfate pentahydrate (on ice). Because of the sensitivity to light of the reagents, the silver enhancement and gold toning steps were performed in a dark room under a red light. Finally, the cells were post-fixed with 1% w/v osmium tetroxide + 1.5% w/v potassium ferricyanide for 1 h, *en bloc* stained with 2% w/v uranyl acetate for 20 min, dehydrated in a gradient of ethanol, embedded in Procure resin, and polymerised at 60°C for 48 h. Polymerised resin blocks were sectioned using an ultramicrotome (Ultracut 7, Leica Microsystems, Wetzlar, Germany) to generate 60 nm ultra-thin sections that were collected on 200 mesh copper grids. Sections were post-stained with 2% w/v uranyl acetate and Reynold’s lead citrate for 10 min, each before imaging under a transmission electron microscope at 200 kV (G2 Tecnai, FEI, Hillsboro, OR, USA).

### Mouse tissue

Tissue collection protocols were approved by the Garvan Institute and St. Vincent’s Hospital Animal Ethics Committee and were in accordance with the guidelines of the Australian code for the care and use of animals for scientific purposes (8th edition, National Health and Medical Research Council, Canberra, ACT, Australia, 2013) and the Guide for the Care and Use of Laboratory Animals (8th edition, National Research Council, USA, 2011). Piezo1-Tdtomato mice (The Jackson Laboratory Stock No: 029214) were housed in light boxes and entrained to a 12:12 light: dark cycle for 1 week before the experiments. Mice aged 10 weeks old were euthanized with carbon dioxide, lung tissue harvested and immediately homogenized in RIPA buffer, as above, for protein extraction.

### RBC collection

Human RBCs were prepared from blood obtained by venipuncture from the cubital fossa of normal informed-consenting donors under approval from the University of Sydney Human Ethics Committee (signed, approved consent form; Project No. 2012/2882). The blood was anticoagulated with 15 IU mL^-1^ porcine-gut heparin (Sigma-Aldrich). The blood was centrifuged in 50 mL Falcon tubes at 3000 × g for 5 min at 4°C to sediment the RBCs, and the plasma and buffy coat were removed by vacuum-pump aspiration. RBCs were lysed prior to blotting in RIPA buffer as documented above.

### Immunofluorescence and Western blotting of unroofed fibroblasts

The unroofing method for selectively isolating basal membranes of cells was adapted from^35^. Briefly, human BJ-5ta-hTERT foreskin fibroblasts were seeded onto glass coverslips pre-coated with 90 nM fibronectin (20 μg mL^-1^; Cat F1141-5MG, Sigma-Aldrich). After 24 h, the cells were washed with PBS then incubated in an hypo-osmotic buffer containing 2.5 mM triethanolamine (TEA) (pH 7.0) for 3 min at room temperature. The cells were seen to be slightly swollen at this stage in the preparation. Immediately after, the TEA medium was removed and the coverslips were washed with PBS containing protease inhibitor (1 tablet of cOmplete Mini, EDTA-free per 10 mL of 1 × PBS) using a fine-tip transfer pipette.

Once the cells were unroofed (the extent of this was assessed under a light microscope: unroofed cells had no nucleus), the coverslips were transferred to 4% v/v PFA in cytoskeleton buffer [CB; 10 mM 2-(*N*-morpholino)ethanesulfonic acid (MES) pH 6.1, 138 mM KCl, 3 mM MgCl_2_, 2 mM EGTA] and left to fix for 10 min. The fixed cells were then labelled for actin, the focal adhesion proteins, and nuclei using, respectively, phalloidin conjugated to AlexaFluor568 (A12380, ThermoScientific), a mouse anti-vinculin antibody (1:200; V9131, Sigma-Aldrich), and DAPI. Images were acquired using a widefield fluorescence microscope at 63 × (Ti2-E, Nikon, Tokyo, Japan). Alternatively, after unroofing the cells they were lysed in fresh RIPA buffer prior to Western blotting.

### Biotinylation

Human BJ-5ta-hTERT foreskin fibroblasts cells were grown to 70-90% confluency on a 60 mm dish before being biotinylated. Cells were washed three times using ice cold PBS and then incubated with biotin buffer (154 mM NaCl, 10 mM HEPES, 3 mM KCL, 1 mM MgCl_2_, 0.1 mM CaCl_2_, 10 mM glucose, pH 7.6) containing 1 mg/mL Sulfo-NHS-Biotin (ThermoFisher Scientific, EZ-Link™, Lot. No. TI266926) for 1 h on ice. The biotin buffer was then washed off and quenched using 5 mL DMEM containing 25 mM HEPES and 0.25 % w/v gelatin for 10 min on ice. Cells were then washed three times using ice cold PBS and solubilized using 1 mL RIPA buffer. Cell lysates were cleared by centrifugation at 13,000 × g at 4 °C for 20 min. 100 μL supernatant was taken and supplemented with 2 % w/v SDS and 0.8 M urea and designated as the “input” sample. The remaining supernatant was incubated overnight at 4 °C with 100 μL Streptavidin-Agarose beads (Sigma-Aldrich, Lot#: SLBR5741V) blocked using 0.5 % w/v bovine serum albumin for 1 h. Beads were collected and washed three times using RIPA buffer in a 0.8 mL centrifuge column (ThermoFisher Scientific, Pierce, Cat. No. 89868) with centrifugation at 500 × g at 4 °C for 1 min. 100 μL flow-through lysate were taken from the first centrifuge and supplemented with 2 % w/v SDS and 0.8 M urea to be the flow-through comparison. For protein elution, beads were loaded with 100 μL 0.9 % w/v NaCl solution supplemented with 2 % SDS and 0.8 M urea, and boiled at 65 °C for 5 min. Then they were centrifuged at 500 × g at 4 °C for 1 min in the column, and the biotinylated eluate was collected. Identical protein amounts input, biotinylated and flow-through samples were loaded for Western blotting then probed with mouse monoclonal anti-Piezo1 antibody (Cat. No. NBP2-75617, Novus Biologicals) and mouse anti-α-actinin antibody (Santa Cruz Biotechnology, Dallas, TX, USA).

### Ca^2+^ imaging

Piezo1^-/-^ HEK293T cells expressing Piezo1 were seeded into a 96 well plate and incubated with 100 μL of 2 μM Fura-2-AM in DMEM for 30 min at 37°C. Ca^2+^ transients were recorded using a 20 × objective mounted on a Nikon Ti2e microscope and illuminated with a CoolLED pE-340 Fura LED light source.

## Results

### N-linked glycosylation of heterologously expressed Piezo1

First, we used Piezo1 fusion proteins to probe Piezo1 expression because there are no specific anti-Piezo1 antibodies; instead we exploited highly specific antibodies to GFP and mCherry. This allowed us to correct for minor variability in transient transfection by using the GFP or mCherry fluorescence intensity of the cell lysate to calculate the relative amounts required to load for Western blotting. Initial trials at probing Piezo1 expression in Piezo1^-/-^ HEK293T cells with Western blotting showed extensive smearing in the lanes. This was corrected by adding a reducing agent during cell lysis, thus preventing protein cross-linking and aggregation. Figure S1A shows gels run using samples without reducing agents in RIPA buffer compared with samples with 5 mM β-mercaptoethanol, 5 mM TCEP, or 10 mM of the alkylating agent NEM, during cell lysis. We found that NEM treated Piezo1 was slightly larger in molecular size due to the large number of cysteines present in Piezo1. Specifically, if all 57 -SH groups were in a reduced state before modification with NEM (0.125 Da) there would have been an increase in size of 57 × 0.125 Da = 7.125 kDa, thus resulting in a significant net increase.

With lysates from HEK cells, Western blots showed two distinct bands (Fig. 1 A, B; fully glycosylated - FG; core glycosylated – CG and UG; unglycosylated) that resembled Piezo1 from native mouse tissue (specifically aorta)^36^. To determine if the two bands resulted from different extents of glycosylation, Piezo1-GFP lysates were incubated with PNGaseF that only hydrolyses N-linked glycosyl units from the free end of an oligosaccharide chain, and a mixture of deglycosylases that hydrolyse N- and O-linked oligosaccharides. Figure 1B shows the upper band (labelled fully glycosylated - FG) was absent after treatment, while the lower band had increased mobility suggesting reduced size. The molecular correlates of the UG, CG and FG bands are illustrated pictorially in Fig. 1C. PNGaseF alone and the mixture of deglycosylases produced the same effect. The estimated size difference on the removal of N-linked oligosaccharides, based on the calibrating ladder on the gel, was 20 - 30 kDa.

**Figure 1.**
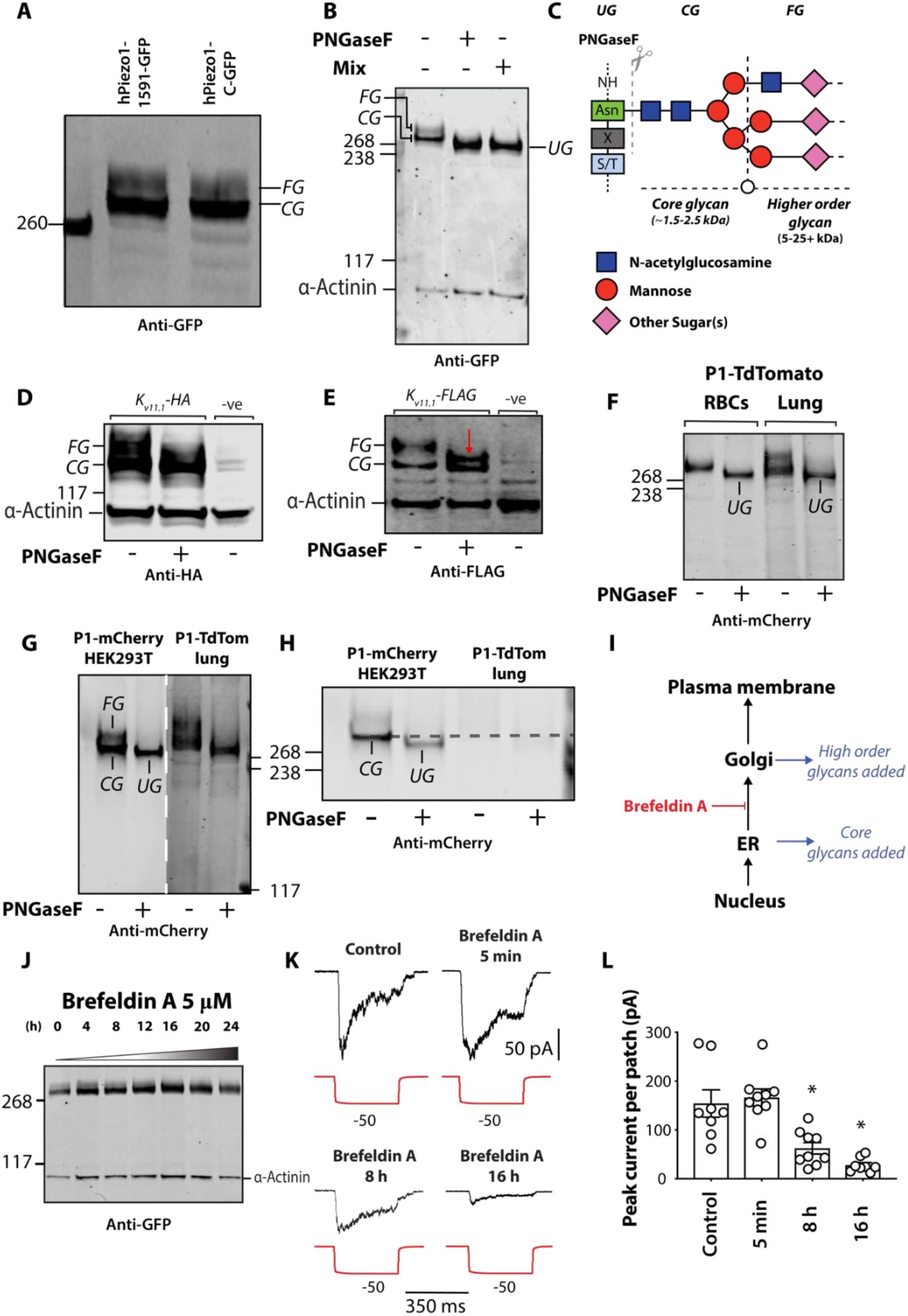
N-linked glycosylation of Piezo1 in a heterologous expression system and in native cell types. (A) A representative Western blot of hPiezo1-1591-GFP and Piezo1-C-GFP expressed in HEK293 cells. (B) A representative Western blot illustrating cleavage of the upper FG (fully glycosylated) band of the Piezo1 doublet in the presence of the enzyme PNGaseF and a mix of O- and N-linked deglycosylase (CG – core-glycosylated, UG – unglycosylated). (C) Schematic illustration of PNGaseF mediated cleavage of N-linked glycans at the molecular level. (D) Representative Western blot showing the effect of PNGaseF digestion on K_v11.1_-HA and (E) K_v11.1_-FLAG protein expressed in HEK293 cells. The red arrow denotes a previously identified PNGaseF resistant K_v11.1_ component. (F) Representative Western blot comparing the effect of PNGaseF digestion of mouse Piezo1-TdTomato from red blood cells (RBCs) and lung tissue. (G) Comparison of Western blot of human Piezo1-1591-mCherry expressed in HEK293 cells with mouse Piezo1-TdTomato from mouse lung tissue. (H) Reduced intensity, of representative blot shown in panel G to illustrate the reduction in size of the lower CG band in addition to the loss of the upper FG band. (I) Schematic representation of where N-glycans are added and the site of action of the fungal metabolite brefeldin A. (J) Representative Western blot of Brefeldin A treatment (0-24 h) on human Piezo1-GFP lysate. (K) Raw electrophysiological traces from cell-attached patches of Brefeldin A treated Piezo1-/- HEK293T expressing Piezo1-GFP recorded at −65 mV. (L) Quantification of peak Piezo1 currents per patch elicited by negative pressure pulses in the presence of brefeldin A for escalating durations. Data expressed as mean ± SEM * p<0.05 determined by Kruskal-Wallis test with Dunn’s post-hoc test.

As a positive control for PNGaseF digestion we used K_v11.1_ the hERG potassium channel. This channel is known to undergo N-linked glycosylation and migrates as two species (Fig. 1D-E), a core glycosylated protein (CG, ~135 kDa) and a fully glycosylated protein (FG) of higher molecular size (~155 kDa)^18^. On treatment with PNGaseF we also saw cleavage of the upper band regardless of which tagged version of the protein was used. This included the generation of a previously reported PNGaseF resistant modification shown in Fig. 1E, labelled with a red arrow.

### N-glycosylation of endogenous Piezo1

The next experiments addressed the question of N-linked glycosylation in cell types that natively express Piezo1. Here, we studied RBCs and lung tissue that are known to have high levels of expression of Piezo1^1^, isolated from a Piezo1-TdTomato reporter mouse. Figure 1F shows that in both RBCs and lysate from lung tissue that the molecular size that Piezo1 runs at is reduced by treatment with PNGaseF. Furthermore, the extent of glycosylation was not altered by the TdTomato tag.

We compared human Piezo1-1591-mCherry to mouse Piezo1-TdTomato extracted from lung tissue that had different tags fused at different positions (Fig. 1F-G). While the sample of mPiezo1-TdTomato fusion protein was larger (~320 kDa), it still clearly contained the species of larger molecular size (glycosylated version), which was reduced in size in the sample that had been treated with PNGaseF. In comparison to the extracts from HEK cells and lung tissue, only one band was observed in the RBC lysate; it became smaller on PNGaseF treatment (Fig. 1F).

To explore further where in the cell the higher order glycans (that we now refer to as “full” glycosylation) were added to Piezo1, the trafficking between the ER and Golgi was inhibited with brefeldin A (Fig. 1I). This fungal metabolite prevents movement of proteins from the ER to the Golgi by disassembling transport vesicles^37^. Treatment of cells with brefeldin A led to the accumulation of the lower band on Western blots, and after 12-16 h the upper fully glycosylated version was no longer evident (Fig. 1J). Therefore, we concluded that the higher order glycans were added to Piezo1 in the Golgi.

If, like K_v11.1_, the fully glycosylated form of Piezo1 was indicative of the mature membrane protein, then brefeldin A treated (>12-16 h) cells should have reduced stretch-activated currents. Indeed, the peak stretch-activated currents recorded from Piezo1^-/-^ HEK293T cells heterologously expressing Piezo1-GFP, compared to those from untreated or acutely treated cells were reduced (Fig. 1K-L). The average current per patch for Piezo1-GFP expressed in Piezo1^-/-^ HEK293T was 154 ± 28 pA (*n* = 8), and this was reduced ~6 fold to 27 ± 5 pA (*n* = 8) after 16 h treatment with brefeldin A. This was not due to blockade of Piezo1 as a 5 min acute treatment with brefeldin A did not affect Piezo1 currents, 164 ± 22 pA (n = 7) (Fig 1K-L).

While probing tagged versions of Piezo1 we identified the same double band in the untagged Piezo1, using a newer mouse monoclonal anti-Piezo1 antibody (Cat. No. NBP2-75617, Novus Biologicals). This antibody reproducibly recognized human Piezo1 which presented as two bands on a Western blot. Importantly, un-transfected Piezo1^-/-^ HEK293T cells served as a negative control (SI Fig. 1B). This antibody also recognized native Piezo1 in human RBCs but failed to recognize mouse Piezo1-GFP recombinantly expressed in Piezo1^-/-^ HEK293T cells. This finding was confirmed when the same Western blot was probed with a GFP antibody (SI Fig. 1B). Hence, we concluded that the untagged protein also underwent glycosylation and that a tag does not influence N-glycosylation.

Endogenous Piezo1 in immortalized human fibroblasts also showed two bands on a Western blot (Fig. 2A). The higher molecular weight band was much more intense than that of human Piezo1 that had been transiently expressed in HEK293 or Neuro2A cells (Figs. 2A-B and SI Fig. 2). The ratio of the upper band compared to the lower band was >10 fold higher in fibroblasts (Figs. 2B). The upper band was assigned to the N-glycosylated protein since treatment with PNGaseF decreased the molecular size, giving coalescence of both bands (Fig. 2A).

**Figure 2.**
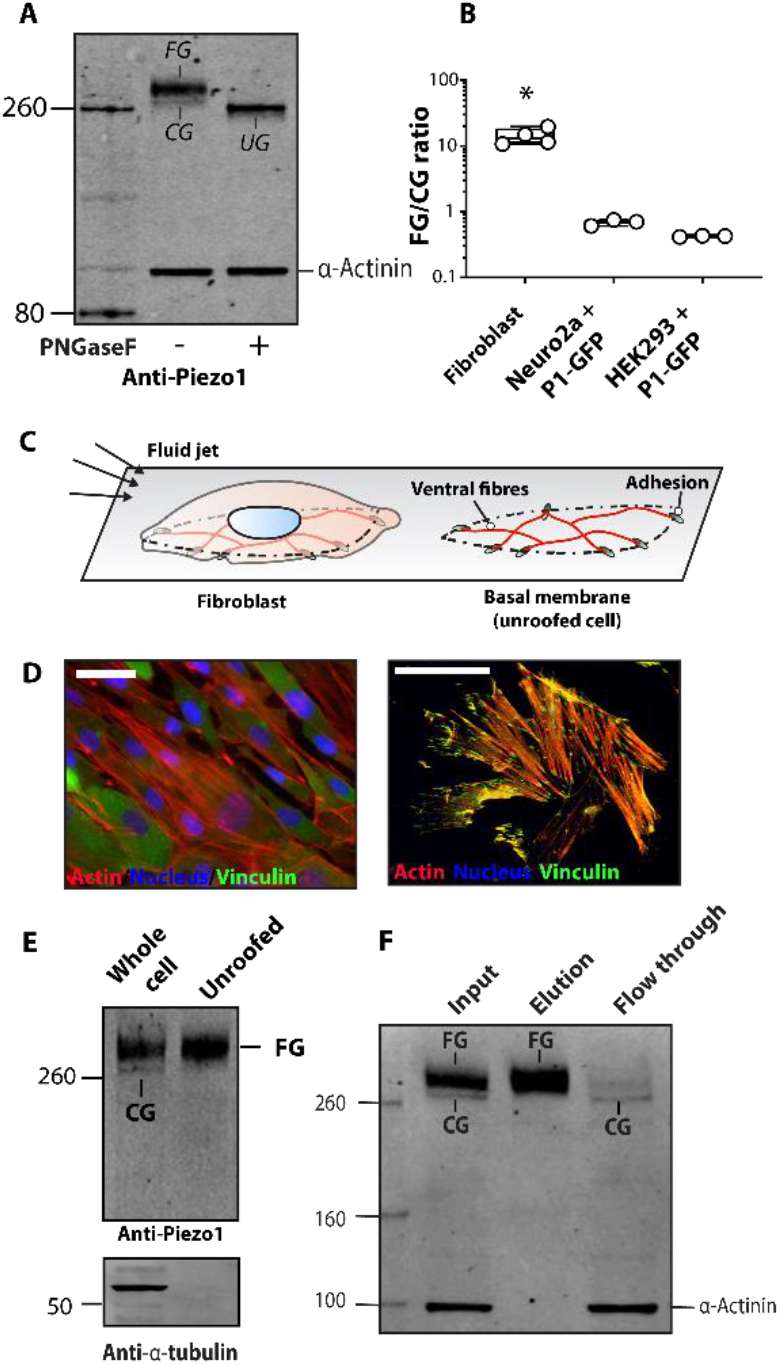
N-glycosylation of endogenous Piezo1 in human fibroblasts. (A) A representative Western blot of untreated immortalized human foreskin fibroblasts versus PNGaseF treated fibroblast lysate probed using the anti-Piezo1 antibody and anti-α-actinin antibody as a loading control. (B) The ratio [FG/CG] of the upper band (FG) and lower band (CG) of fibroblast Piezo1 and Piezo1 heterologously expressed in Neuro2A and HEK293T. (C) Schematic illustration of an intact fibroblast, and an unroofed fibroblast. (D) A representative image of an intact fibroblast (*left panel*), and an unroofed fibroblast (*right panel*) using standard wide-field microscopy and a 63x oil objective. (E) A representative Western blot of intact fibroblast lysate versus unroofed fibroblast lysate probed using anti-Piezo1 antibody and anti α-tubulin antibody as a loading control to confirm unroofing. A representative Western blot of biotinylated immortalized human foreskin fibroblasts. (CG – core-glycosyalted, FG – fully glycosylated) * p<0.05 determined by Kruskal-Wallis test with Dunn’s post-hoc test.

We sought to determine whether the upper band, assigned as the N-glycosylated protein, represented the fully mature plasma membrane protein. For this we used a method to “unroof” fibroblasts. This protocol removes apical plasma membranes, nucleus, and organelles and leaves only the basal membrane attached to the tissue culture substrate (Fig. 2C). The procedure has been widely used in conjunction with various imaging methods^38,39^ and also proteomics^35^ but it has rarely been used for studying plasma membrane localization.

Proof of successful unroofing was obtained by staining for the focal adhesion protein vinculin, and imaging with epifluorescence microscopy (Fig. 2D). While focal adhesions containing vinculin were not clearly defined with intact cells, focal adhesions labelled with vinculin became well defined once the cells were unroofed. We and others have shown that Piezo1 is present in the basal membrane of fibroblasts^40,41^ so Western blots were performed on extracts from intact and unroofed cells. From unroofed cells, only the upper band, which was assigned to the glycosylated protein remained (Fig. 2E); and the unroofed cells also lacked α-tubulin as previously reported^35^. We concluded that the mature membrane pool of Piezo1 in fibroblasts was highly N-glycosylated. This was supported by experiments illustrating only the FG version was biotinylated (Fig. 2F). Furthermore, these experiments confirmed that endogenous Piezo1 in human fibroblasts underwent N-linked glycosylation in a manner similar to Piezo1 that was transiently expressed in HEK cells, Neuro2a cells, and endogenous Piezo1 in mouse lung tissue. Finally, unlike heterologous systems where large amounts of protein are produced, the fully N-glycosylated Piezo1 was the larger proportion of natively expressed Piezo1 in fibroblasts.

### Piezo1 N-glycosylation of two Asn residues in the cap

Having established the existence of N-glycosylation of Piezo1 we sought specification of the sites of this post-translational modification. The on-line program NetNglyc1.0^42^ predicted nine sites that could be N-linked, six in the N-terminus and three in the C-terminus. The three asparagines identified by NetNglyc1.0 in the C-terminal domain are in the cap or C-terminal extracellular domain (CED)^31,32^, one of which is buried and seems unlikely to accept N-glycosylation, while the other two are freely accessible from the extracellular space (Fig. 3A). Previous mass spectrometry identified a glycosylated peptide corresponding to one of these predicted asparagines (N2331)^43^. Both asparagines are highly conserved in homologues of Piezo1 and have the classical signature sequence Asn-Xaa-Ser/Thr for N-linked glycosylation (Fig. 3B). In contrast, these Asn residues are not conserved in Piezo2; but a recent Cryo-Electron Microscopy (Cryo-EM) structure of mouse Piezo2 shows at least one glycan in the cap region^44^. Therefore, to test if the fully glycosylated Piezo1 protein was dependent on either of these asparagines, we created single mutants (N2294Q, N2331Q), and a double mutant (N2294Q and N2331Q; also called “CapQQ”).

**Figure 3.**
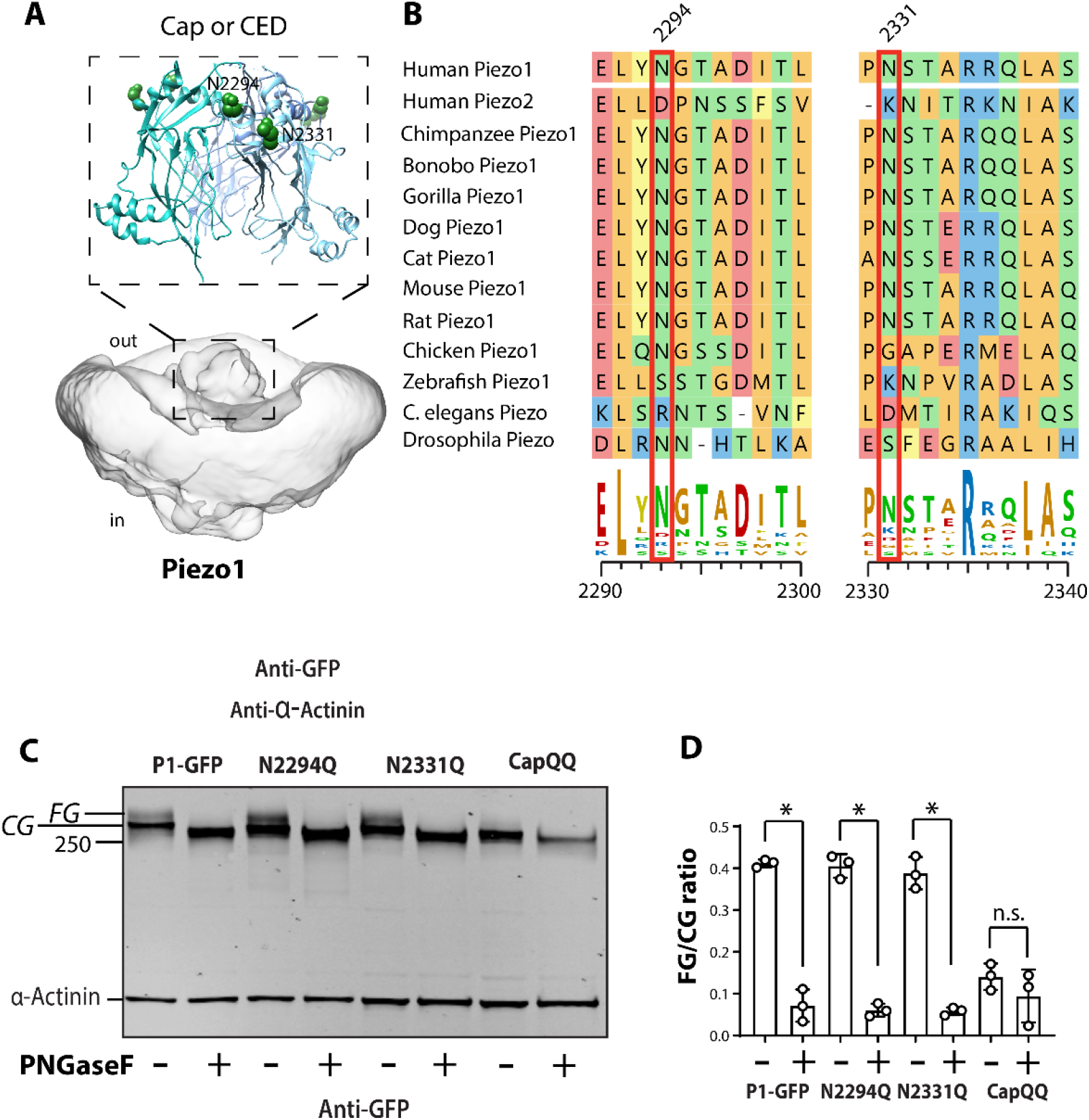
Influence of Asparagine residues in the cap region of human Piezo1 on N-Glycosylation. (A) Two asparagine residues present in the Cap domain of Piezo1. (B) Multiple sequence alignment illustrates these asparagine residues are highly conserved in Piezo1 homologues. (C) Representative blot showing the effect of PNGaseF treatment on lysate from HEK293 cells expressing Piezo1-GFP, N2294. N2331Q and N2294Q/N2331Q (CapQQ). (D) Quantification of the ratio [FG/CG] of the upper band (FG) and lower band (CG) with and without PNGaseF for Asn mutants in the Cap region. * p<0.05 determined by Kruskal-Wallis test with Dunn’s post-hoc test.

Only a minor decrease occurred in the amount (as measured by Western blots) of the fully glycosylated species in the single mutants but there was complete abolition of the upper band in the double mutant (N2294Q and N2331Q) (Fig. 3C), which mirrored the effect of treatment with PNGaseF.

We compared the intensity of the upper fully glycosylated (FG) band with the lower core glycosylated (CG) band in replicate experiments (Fig. 3D). Removal of the oligosaccharides from both the single Piezo1 mutants and control WT-Piezo1-GFP treated with PNGaseF are shown in Figs. 3E-F. Comparison between PNGaseF treatment of WT, N2294Q, N2331Q showed patterns that suggested that the upper FG band was removed and the size of the lower bands were reduced to the level of the un-glycosylated form of the protein. In addition, Western blots of the protein from the double mutant N2294Q/N2331Q showed no upper (fully glycosylated) band while the lower core glycosylated protein was reduced in size (Fig. 3E). This finding suggested that core glycosylation also occurred in the N-terminal domain of Piezo1. This aligns with published findings on Piezo2 that show glycans in the propeller region^44^.

The FG form of an ion channel often indicates the functional form, so we tested if changes in Western blots correlated with electrophysiological analysis. A high-speed pressure clamp was used to apply negative pressure to cell-attached patches of transiently transfected Piezo1^-/-^ HEK293T cells. The two single mutants showed normal stretch-activated responses (Fig. 4A-B), which was consistent with two bands on Western blots. Given strong evidence that the cap is involved in Piezo1 inactivation^45^, we tested if inactivation was modified, but neither of the Asn to Gln mutants affected inactivation time constants in cell-attached patches (Fig. 4C). However, the N2294Q/N2331Q mutant had almost no stretch-activated current (Fig. 4A-B), which was consistent with the Western blot pattern and indicated that the FG Piezo1 protein represented the membrane protein pool (Fig. 3E).

**Figure 4.**
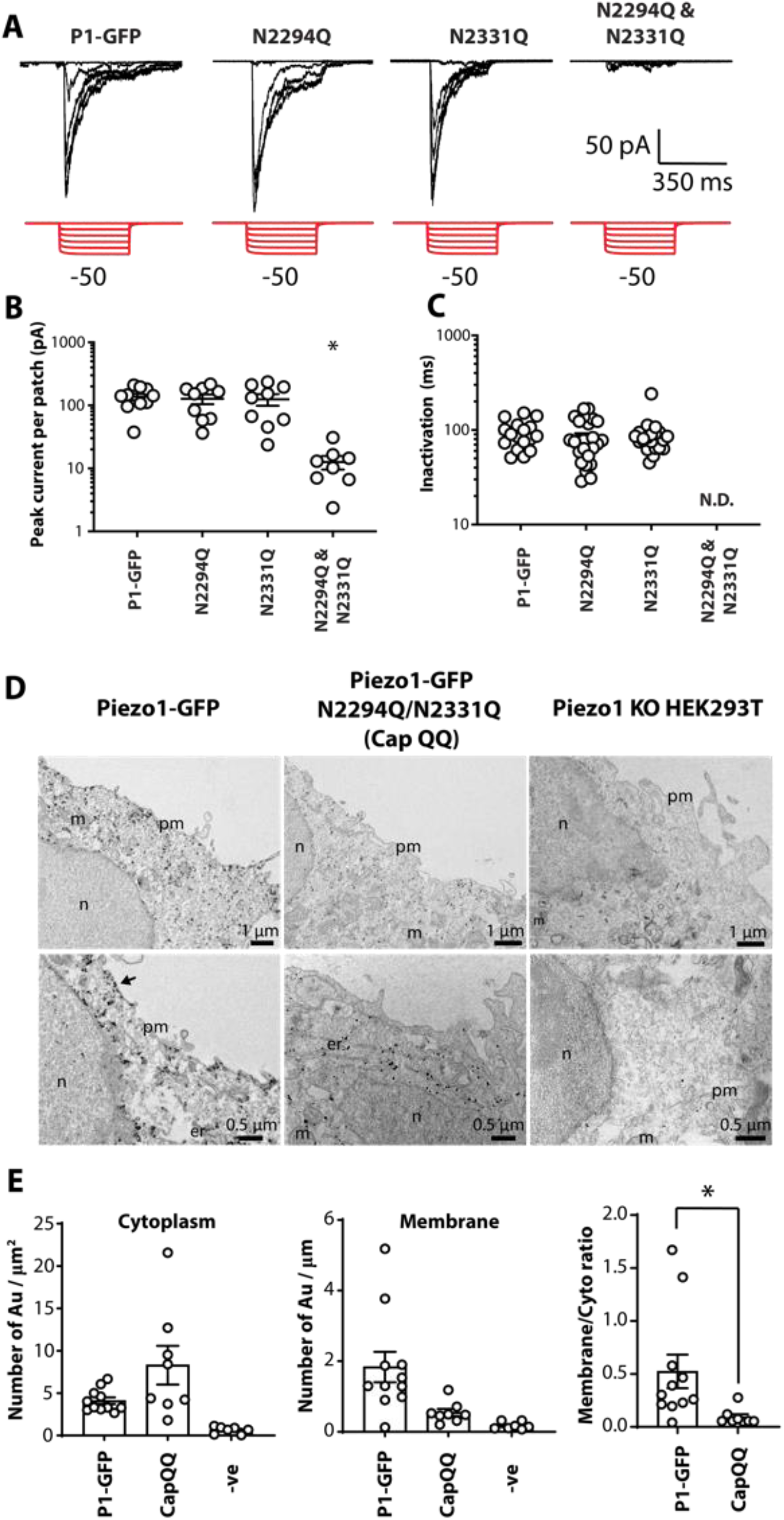
Effect of mutation of N2294 and N2331 on mechanically induced gating of Piezo1. (A) Electrophysiological recordings of HEK293T Piezo1^-/-^ expressing Piezo1-GFP, N2294, N2331Q and N2294Q/N2331Q in the cell-attached configuration in response to negative pressure applied using a high-speed pressure-clamp (red). (B) Quantification of peak current elicited per patch. (C) Quantification of inactivation t of Piezo1-GFP, N2294, N2331Q and N2294Q/N2331Q from cell-attached recordings (ND - not determined). (D) Immunogold labelling using mouse monoclonal anti-Piezo1 primary antibody and electron microscopy of HEK293T Piezo1^-/-^ (KO) and cells expressing Piezo1-GFP and the CapQQ mutant (N2294Q/N2331Q)[pm-plasma membrane, n-nucleus, m-mitochondria, er-endoplasmic reticulum]. (E) Quantification of immunogold labelling in the cytoplasm (per μm^2^), membrane (per μm) and the ratio of membrane to cytoplasmic labelling using immunogold comparing Piezo1 to the CapQQ mutant (N2294Q/N2331Q). *p<0.05 determined using Mann-Whitney U test.

From these data we could not rule out that N-glycosylation may also be required for stretch activation of Piezo1. The processing of N-glycans from high-mannose to higher molecular weight glycans (>10 kDa in size - termed complex or hybrid) requires the enzyme N-acetylglucosaminyl-transferase I (GnT1, also known as MGAT1). Structural biologists have widely used HEK293S GnT1^-/-^ cells to restrict the heterogeneity introduced by N-linked glycosylation when attempting to determine structures using X-ray crystallography, and more recently Cryo-EM^46^. As these cells cannot process higher order glycans (hybrid or complex), we asked two questions: (1) “Is the fully glycosylated species of human Piezo1 present on Western blots from lysates of GnT1^-/-^ cells?” (2) “Can these cells support stretch induced gating of Piezo1 channels?”

First, Piezo1-GFP channels expressed in HEK293S GnT1^-/-^ cells presented as a single band on Western blots consistent with the upper band being a glycosylated species containing higher order glycans (SI Fig 3A). Piezo1-GFP also exhibited stretch-activated currents suggesting higher order glycosylation was not needed for stretch activation. Much like transient transfection in Piezo1^-/-^ HEK293T cells expression of the CapQQ mutant generated negligible stretch-activated currents when transiently transfected in HEK293S GnT1^-/-^ (SI Fig 3B-C). Here we should note that the residual current seen when transfecting the CapQQ (N2294Q/N2331Q) in HEK293S GnT1^-/-^ could have come from the endogenous Piezo1 in this cell type. The sensitivity of the stretch-activated currents of Piezo1-GFP was lower when expressed in HEK293S GnT1^-/-^ cells (SI Fig 3D). The expression of Piezo1-GFP in GnT1^-/-^ cells suggested that higher order glycans are not needed for stretch activation and that only the core-glycans were ultimately necessary for transit through the Golgi to the plasma membrane.

To support the hypothesis that the double mutant (N2294Q/N2331Q) that lacked higher order glycosylation was trafficking-defective we used ratiometric Ca^2+^ imaging to explore the effect of Yoda-1 (2 μM) on Ca^2+^ influx. Fluorescence did not change in cells expressing the double mutant (N2294Q/N2331Q) on adding Yoda-1 thus indicating that the Ca^2+^ concentration did not rise (SI Fig 4). These mutations do not reside near the putative Yoda-1 binding pocket so we surmised that they would be unlikely to influence Yoda-1 binding^47^.

We confirmed the trafficking defect using nano-gold immunolabeling and transmission electron microscopy (TEM) in combination with Piezo1^-/-^ HEK293T cells. The micrographs clearly showed WT Piezo1 had reached the plasma membrane, while the N2294Q/N2331Q double mutant showed little or no membrane labelling (Fig. 4D). Quantification of the ratio of membrane versus intracellular nano-gold staining of Piezo1 and N2294Q/N2331Q showed a marked reduction in membrane labelling of N2294Q/N2331Q, which was consistent with a trafficking defect (Fig. 4E). Nano-gold labelling was not seen in un-transfected Piezo1^-/-^ HEK293T cells.

To corroborate the veracity of the antibody’s specificity, in addition to the Western blots shown in SI Fig1, and immunogold negative controls (Fig. 4D: rightmost panels), immunofluorescence was used in combination with Piezo1-GFP expressed in Piezo1^-/-^ HEK293T cells. SI Fig. 5 shows no staining of Piezo1^-/-^ HEK293T cells and more convincingly that un-transfected cells were not labelled (SI Fig 5).

### Piezo1 higher order N-linked glycosylation in the propellers

To determine the location of N-linked glycosylation in Piezo1 channels use was made of a split human Piezo1 construct generated by the Gottlieb laboratory^48^. This construct has two portions of human Piezo1; the first extends from residue 1 to 1591 fused to mCherry (N-terminal portion - propellers) and the second starts with GFP that is fused to residue 1592 and extends to residue 2521 (C-terminal portion – pore and cap) (Fig. 5A). Western blotting showed that this construct was expressed as two separate molecular entities smaller in size than WT Piezo1 (Fig. 5B).

**Figure 5.**
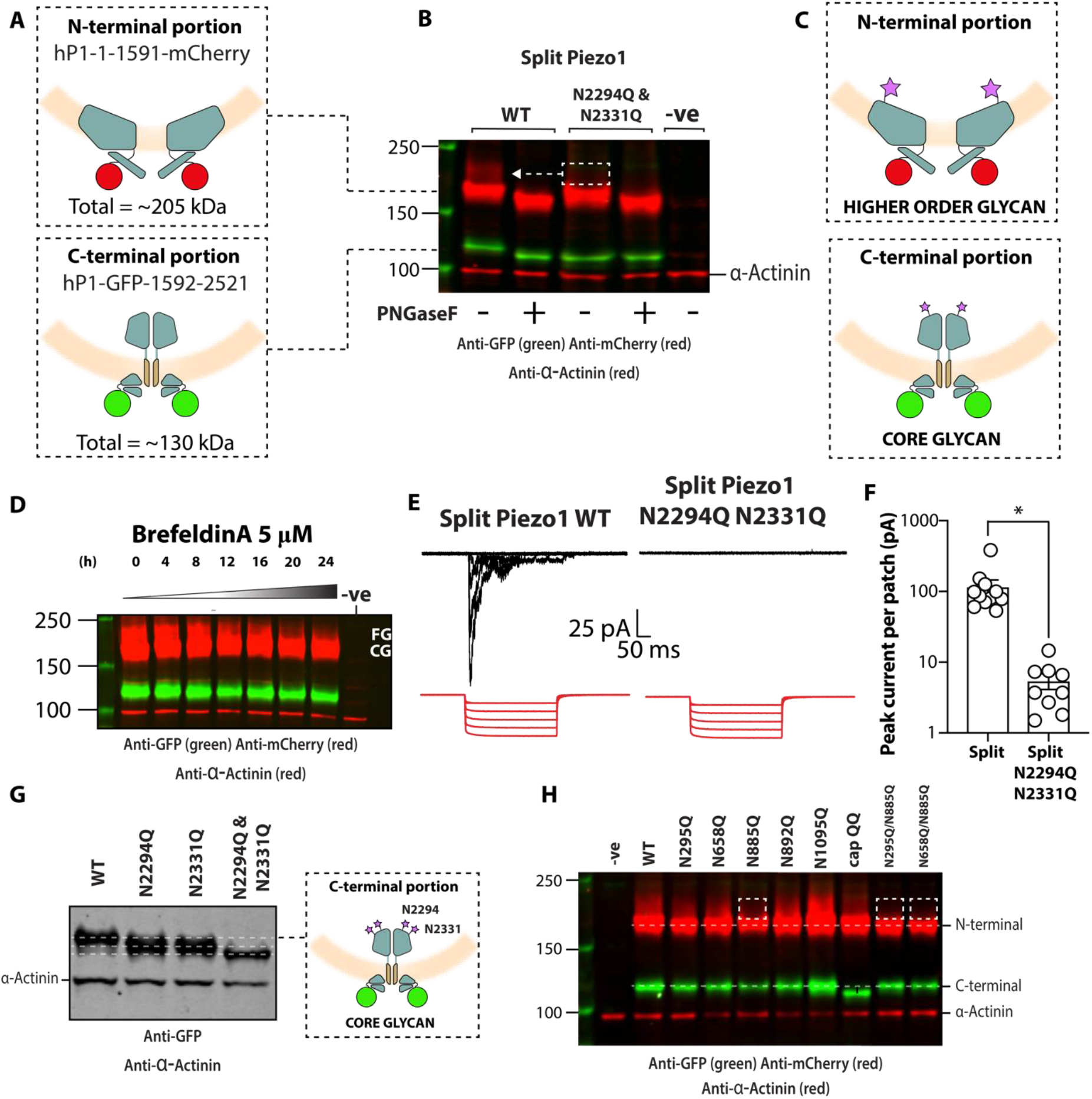
N-glycosylation of Piezo1 in a split construct. (A) Diagram depicting the human split Piezo1 protein. (B) Representative blot showing the split human Piezo1 protein and the split human Piezo1 protein with the double cap mutant N2294Q and N2331Q with and without PNGaseF treatment. (C) Diagram illustrating where glycans are likely to be located. (D) Effect of brefeldin A treatment overtime on the human split Piezo1 protein. (E) Electrophysiological recordings of Piezo1^-/-^ HEK293T cells expressing human split Piezo1 and the double cap mutant N2294Q and N2331Q in the cell-attached configuration in response to negative pressure applied using a high-speed pressure-clamp (red). (F) Quantification of peak current elicited per patch. (G) Representative Western blot of the C-terminal domain of the human Piezo1 split protein compared with single N2294Q and N2331Q and the double cap mutant N2294Q and N2331Q (CapQQ). (H) Representative Western blot showing the comparison of N-terminal Asn to Gln mutations in the N-terminal portion of the human split Piezo1 and the double cap mutant N2294Q and N2331Q (CapQQ). *p<0.05 determined by Mann-Whitney-U test. -ve represents an un-transfected control.

The size of the C-terminal portion of the split protein (Fig. 5B; shown in green) was minimally affected by PNGaseF treatment (~5 kDa), the size of which was indicative of 1-2 core glycans (~ 2-5 kDa). Instead, the larger glycan appeared to be present on the N-terminal portion of the protein (Fig. 5B-C). The N-terminal portion (colored red) migrated as two bands. The larger more diffuse band was abolished and the lower band reduced in size on treatment with PNGaseF (Fig. 5B).

The N2294Q/N2331Q double mutant of the split construct gave a C-terminal domain that was unaffected by PNGaseF treatment, again indicating that it was un-glycosylated; but the N-terminal portion had no higher order glycosylation (Fig. 5B, white box). Thus, the two Asn residues in the cap appeared to determine the glycosylation status of the propeller asparagines. The higher order glycosylation in the N-terminal fragment was also ablated by incubation with brefeldin A (Fig. 5D). This is consistent with the split protein being processed in a similar fashion to the full-length Piezo1 (Fig. 1I).

The WT split protein produced stretch-activated current when expressed in Piezo1^-/-^ HEK293T cells, while the double mutant (N2294Q and N2331Q) did not (Fig. 5E-F). As a means of providing further supporting evidence for core glycans being added to Piezo1 at both N2294 and N2331, we made single mutants of the same split construct and compared the molecular size to the split WT and double mutant (N2294Q and N2331Q). The corresponding blot showed a similar size between the single mutants (Fig. 5G). The size was less than that of WT but larger than the double mutant, further suggesting that both asparagines were glycosylated (Fig. 5G).

The split construct that produced smaller Piezo1 protein fragments was used to determine if the propeller domains contained a site for higher order glycosylation. There are six residues that are possibilities to undergo N-linked glycosylation in the N-terminus of human Piezo1: N295, N658, N885, N892, N1095 and N1222. Of these, five out of the six are in extracellular loops while one (N1222) is part of a transmembrane helix, so it was not explored further. Single Asn to Gln mutations were made to prevent N-linked glycosylation in the split construct. Again, the CapQQ (N2294Q and N2331Q) mutant had a smaller C-terminal fragment that was concluded to be due to a lack of core-glycans in the cap; and N885Q had limited higher order glycosylation on their N-terminal fragments despite containing the two critical Asn residues in the cap (Fig. 5H).

Subsequently, constructs of full-length Piezo1 were made in which all five Asn residues were mutated to Gln. Stretch activated currents were measured for each of these full length Piezo1 variants (Fig. 6A). Figure 6B shows the peak currents of cell-attached patches when these mutants were expressed in Piezo1^-/-^ HEK293T cells. Specifically, there was a large reduction in stretch activated currents in N885Q; and Western blotting confirmed an almost complete loss of higher order glycosylation in this single mutant (Fig. 6C). None of the five mutations altered unitary conductance from WT (47 ± 2 pS; n = 5) (SI Fig. 6).

**Figure 6.**
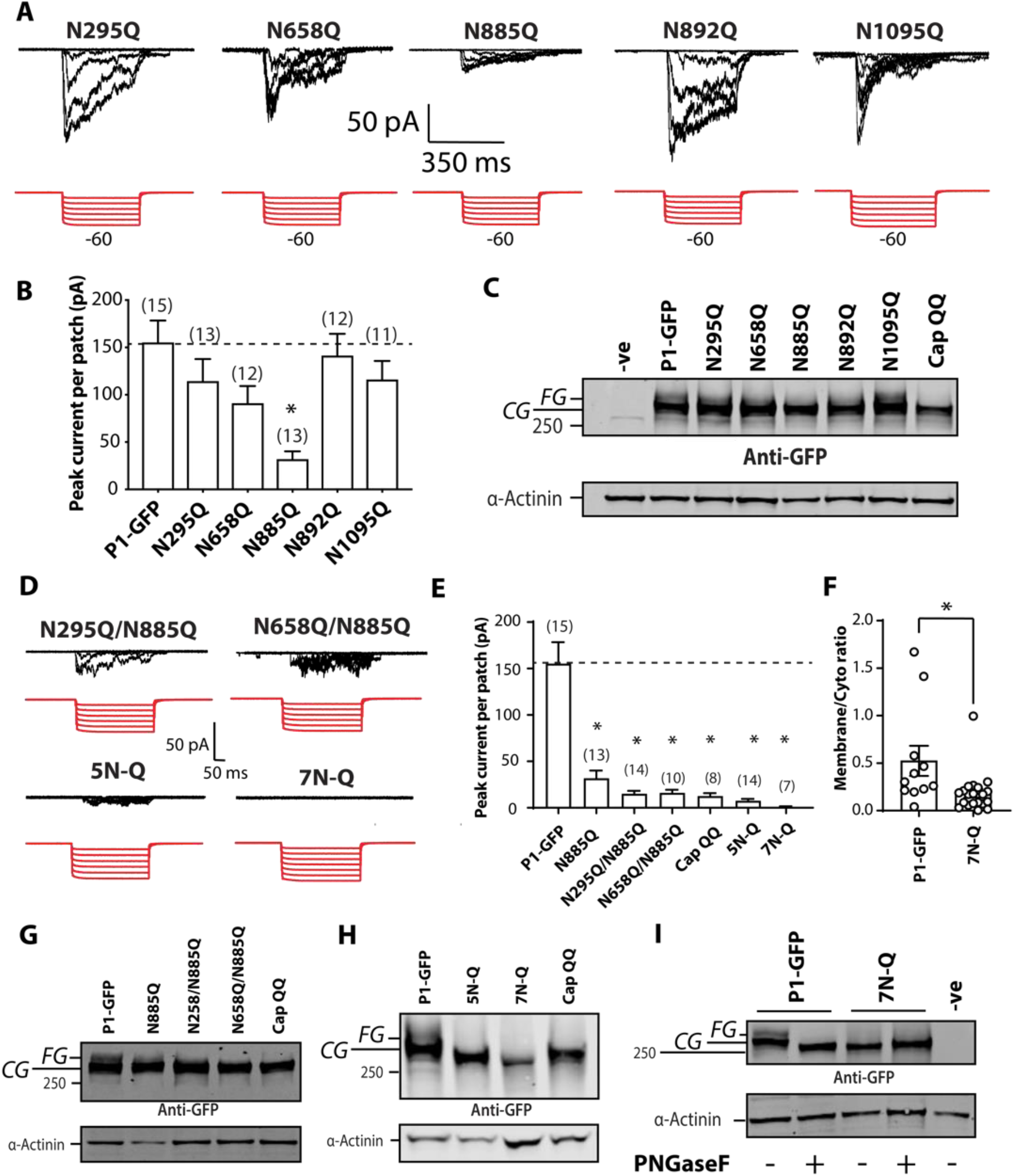
Higher order N-glycosylation in the N-terminus of Piezo1. (A) Electrophysiological recordings of Piezo1^-/-^ HEK293T expressing single Asn to Gln Piezo1-GFP mutations in the N-terminus (N295Q, N658Q, N885Q, N892Q, N1095Q) in the cell-attached configuration in response to negative pressure applied using a high-speed pressure-clamp (red). (B) Quantification of peak current elicited per patch for point mutations shown in A compared to current from Piezo1-GFP. (C) Representative Western blot of all Asn to Gln mutants in the N-terminus of Piezo1 compared to the double mutant in the Cap region (CapQQ). (D) Electrophysiological recordings of HEK293T Piezo1^-/-^ expressing Piezo1 with multiple Asn to Gln mutations (as indicated) in the cell-attached configuration in response to negative pressure applied using a high-speed pressure-clamp (red). [5N-Q - N295Q, N658Q, N885Q, N892Q, N1095Q and 7N-Q - N295Q, N658Q, N885Q, N892Q, N1095Q, N2294Q, N2331Q]. (E) Quantification of peak current elicited per patch for mutation combinations shown in D compared to current from Piezo1-GFP, N885Q and CapQQ. (F) Membrane to cytoplasmic ratio of immunogold labelling of the 7N-Q mutation compared to human Piezo1. (G) Representative Western blot showing the comparison between Asn to Gln mutations of single and double mutations illustrating the strong impact of N885 mutation on the upper FG band of Piezo1 heterologously expressed in Piezo1^-/-^ HEK293T. (H) Representative blot showing the comparison between multiple Asn to Gln mutations 5N-Q, 7N-Q and the CapQQ (N2294Q/N2331Q). (I) Representative blot showing the comparison between PNGaseF digested WT and the 7N-Q mutant. * p<0.05 determined by Kruskal-Wallis test with Dunn’s post-hoc test or Mann-Whitney-U test. -ve represents an un-transfected control.

We noted that the higher order glycosylation of N658Q was different as the patch clamp data showed a mild reduction in stretch-activated currents (Fig 6C). Based on these findings combinations of Asn to Gln mutants were constructed.

A double mutant of N295Q and N885Q gave stretch-activated responses that largely followed the current generated from the single N885Q mutant; while a combination of N658Q and N885Q further reduced the stretch activated current (Fig. 6D-E). Combining all five N-terminal Asn residues to Gln mutants (N295Q, N658Q, N885Q, N892Q, N1095Q), or all seven Asn residues subjected to electrophysiological analysis in this study (N295Q, N658Q, N885Q, N892Q, N1095Q, N2294Q, N2331Q) resulted in maximum currents that fell to almost zero in response to negative pressure pulses (Fig. 6E). The construct harboring seven Asn to Gln (7N-Q) mutations did not reach the plasma membrane as assessed using immunogold labelling in a manner similar to that seen with the CapQQ (N2294Q/N2331Q) in Figure 3.

Western blot analysis of the double mutants, particularly N658Q and N885Q showed an almost complete loss of the higher order glycosylation in a manner similar to the CapQQ (N2294Q/N2331Q) (Fig. 6G). The 5N-Q (N295Q, N658Q, N885Q, N892Q, N1095Q) and 7N-Q (N295Q, N658Q, N885Q, N892Q, N1095Q, N2294Q, N2331Q) mutants on Western blots, had bands that were consistent with full-length protein but there was a complete lack of the upper fully glycosylated species (Fig. 6H). Moreover, the lower band in both the 5N-Q and 7N-Q Piezo1 mutants were smaller than that of the core-glycosylated Piezo1-GFP. This finding was consistent with the lack of core-glycans being attached at these sites. As final evidence that all the N-linked glycans were removed from the 7N-Q mutant, we treated it with PNGaseF and saw that it had no effect on protein size (Fig. 6H).

### Lack of higher order N-glycosylation in trafficking defective Piezo1 mutants

As the data above suggested that the higher order glycosylation was a surrogate for normal membrane trafficking in human Piezo1 channels, we investigated if disease-linked variants were identifiable by using this approach. Such an approach has been extensively used in studies of loss-of-function disease-causing variants in K_v11.1_ that as mentioned previously cause LQTS2^18^. First, variants linked to generalized lymphatic dysplasia (L939M^27^ and G2029R^28^) were studied.

Much like the Cap QQ (N2294Q/N2331Q) mutant, G2029R shows no higher order glycosylation (Fig. 7A). In comparison, L939M Western blots appeared similar to those of the WT, with two bands. The relative amounts were quantified using the intensity of the upper band (fully glycosylated, FG) versus that of the lower band (core glycosylated, CG) and these showed a marked reduction of the FG band in the Cap QQ (N2294Q/N2331Q) and the G2029R mutants (Fig. 7B). The latter mutant had already been convincingly shown with immunofluorescence to be trafficking-defective^28^, a fact we confirmed here using immunogold electron microscopy (Fig. 7C). Supporting this finding, little to no stretch-activated current was seen in cell-attached patches expressing G2029R in comparison to L939M in which peak currents were comparable to WT (Fig. 7D-E). This finding provided further support for the idea that the presence of higher order N-linked glycosylation is indicative of normal membrane trafficking.

**Figure 7.**
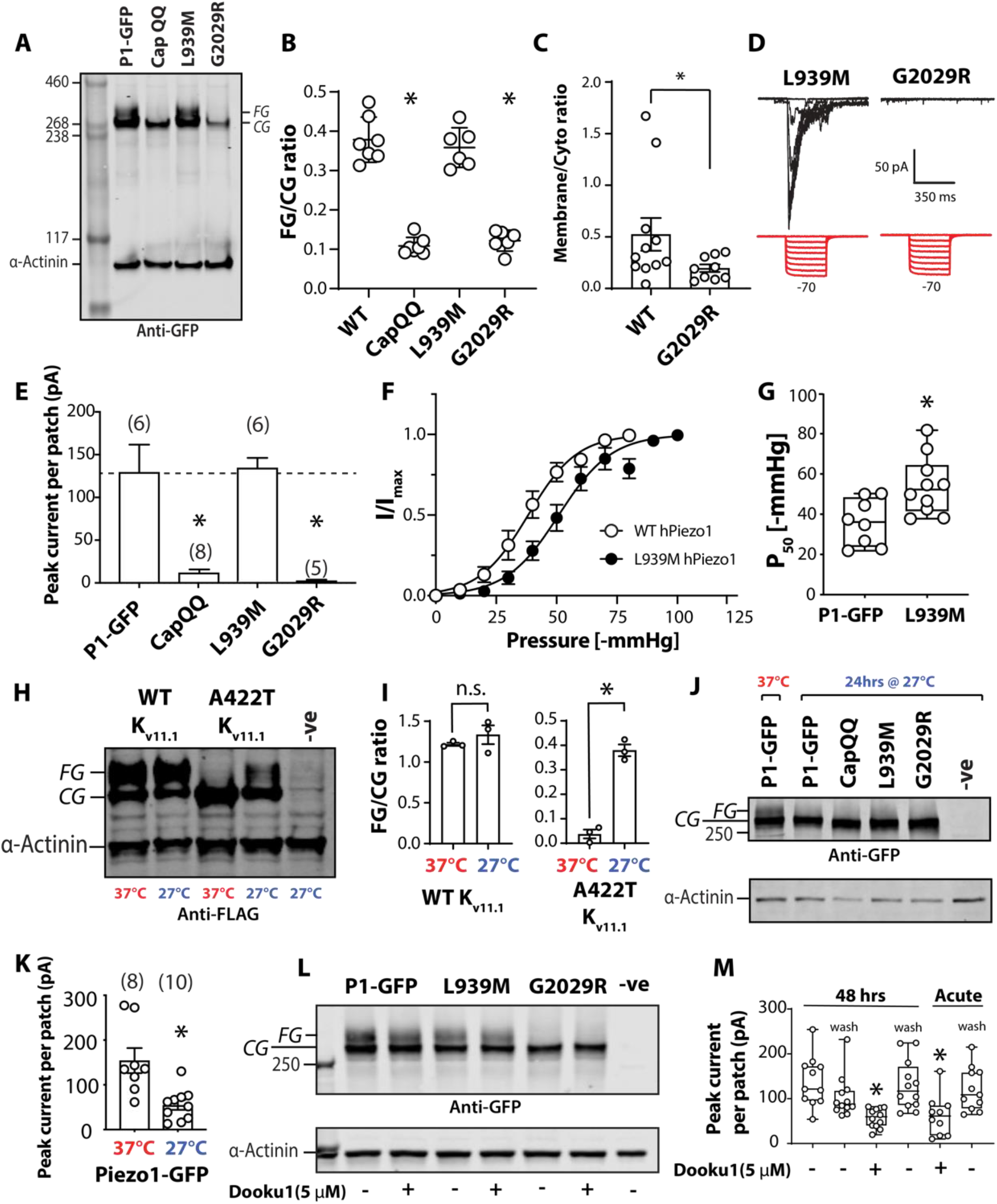
N-glycosylation status of Piezo1 variants linked to generalized lymphatic dysplasia (GLD). (A) Representative Western blot showing Piezo1 protein, CapQQ (N2294Q/N2331Q) and two GLD associated mutations (L939M and G2029R). (B) Relative quantification of intensity of upper band (FG) over lower band (CG) for Piezo1, CapQQ, L939M and G2029R. (C) Membrane to cytoplasmic ratio of immunogold of G2029R compared to human Piezo1. (D) Electrophysiological recordings of Piezo1^-/-^ HEK293T expressing L939M and G2029R in the cell-attached configuration in response to negative pressure applied using a high-speed pressure-clamp (red). (E) Quantification of peak current elicited per patch for point mutations shown in D compared to current from Piezo1-GFP and CapQQ mutant. (F) Pressure response curve for Piezo1-GFP against L939M expressed in Piezo1^-/-^ HEK293T. (G) Box and whiskers plot showing the P50 [-mmHg] of Piezo1-GFP and L939M variant. (H) Representative Western blot showing that 27°C treatment for 24 hours does not change the upper FG band of K_v11.1_ WT markedly, but considerably increases the upper FG band while concomitantly decreases the lower CG band of K_v11.1_ A422T mutant which is temperature rescuable. (I) Quantification of upper FG band/lower CG band ratio of samples shown in H. (J) Representative Western blot including HEK293T cells incubated at 37°C while expressing Piezo1-GFP as a control, and 27°C for 24 hours for Piezo1-GFP, CapQQ, L939M, and G2029R. (K) Quantification of peak current elicited per patch for Piezo1^-/-^ HEK293T cells expressing P1-GFP incubated at 37°C and 27°C for 24 hours. (L) Representative Western blot comparing the effect of 5 μM Dooku1 treatment for 48 h on upper FG bands of P1-GFP, L939M and G2029R. (M) Quantification of peak current elicited per patch showing the effect of treatment with 5 μM Dooku1 on P1-GFP. This includes both 48 h of treatment with Dooku1 compared to washout and acute treatment compared to washout. * p<0.05 determined by Kruskal-Wallis test with Dunn’s post-hoc test or Mann-Whitney-U test. -ve represents an un-transfected control.

While the maximum stretch-activated L939M currents were comparable to WT, they did show a modest rightward shift in the pressure-response curve (Fig. 7F-G). It is important to note that we were not aiming to definitively ascribe disease causation to L939M, in fact, the patient with this mutation also had other missense variants reported in Piezo1 (F2458L, R2456C), which we did not test^27^. What was evident was that WT and L939M Piezo1 both showed a double band on a Western blot, and they gave stretch activated currents; and both CapQQ and G2029R only had the lower band in the Western blots and gave limited stretch activated currents.

Given that we could clearly distinguish a trafficking defective human Piezo1 variant (G2029R) from WT or a variant that reaches the plasma membrane (L939M), we posit that this experiment could serve as an assay for ameliorated trafficking.

### Temperature effects on trafficking

Again, using parallels with the K_v11.1_ and CFTR literature, we attempted to rescue aberrant trafficking using two methods. The first was low temperature treatment, and the second was a pharmacological approach^18,22,49,50^. Specifically, a cohort of K_v11.1_ and CFTR variants could be rescued at low temperature (<30 °C) which is thought to improve protein folding and hence trafficking. The same experimental protocol was used for the Piezo1 variants, expressed in HEK293 cells for 24 h at 27 °C. WT K_v11.1_ was used with a temperature rescuable mutant A422T as a positive control. 24 h at 27 °C increased the amount of fully glycosylated K_v11.1_ A422T which was consistent with previous reports (Fig. 7H-I). With the same protocol for Piezo1the density of the upper band in the Western blot of the G2029R mutant was not increased. Furthermore, the treatment reduced the upper band of WT and L939M suggesting amelioration of trafficking (Fig. 7J). The lower membrane expression level was confirmed by using patch clamp experiments (Fig. 7K). Importantly, all patch clamp experiments were carried out at room temperature as previous reports suggested that Piezo1 activity is temperature dependent^51^. These findings provided further evidence that the intensity of the upper band in Western blots of the HEK Piezo1 proteins reported on channel trafficking to the plasma membrane.

### Drug effects on trafficking

K_v11.1_ is stabilized by the channel blocker E4031, which improves membrane trafficking^22^. So we tested if the antagonist of Yoda-1 activation Dooku-1^52^ offers the same type of chaperone effect on Piezo1. Also, if Dooku-1 did not inhibit stretch-activation of Piezo1, this type of molecule might be used therapeutically if it could improve trafficking.

Treatment of HEK cells for 48 h with 5 μM Dooku-1 (higher concentrations could not be attained due to its low solubility) did not improve the intensity of the upper Western blot FG band of the WT, L939M or the G2029R mutant of Piezo1 (Fig. 7L). In the process of confirming this outcome with patch clamp analysis, the current per patch of Piezo1 was seen to be reduced by 30-60% (Fig. 7M). After washout, peak currents per patch returned to >80% of untreated levels. This suggested that Dooku-1 perturbed Piezo1 stretch-activation. Therefore, we tested acute addition of 5 μM Dooku-1. Indeed, it reduced stretch activation by ~50%, which was largely reversed on washout.

This warrants further study. While neither temperature nor Dooku1 could reverse amelioration of trafficking this does provide the basis for future studies to look at a wider range of loss-of-function Piezo1 mutants. However, it was first necessary to see if modified N-glycosylation was applicable to other reported disease-linked Piezo1 variants.

### Co-expression of trafficking defective Piezo1 mutants

Heterozygous Piezo1 missense variants have been linked to disease. To broaden the relevance our data, we investigated more variants with genetic loci that are linked to bicuspid aortic valve (BAV)^29^ and examined how co-expression of disease-linked variants with WT Piezo1 might affect Piezo1 activity. First, WT Piezo1-GFP was co-expressed with equivalent amounts of DNA of L939M and G2029R. The stretch activated currents elicited by negative pressure application from a high-speed pressure clamp (Fig. 8A) were recorded followed by the variants at loci linked to BAV (Fig. 8B-D). Co-expression of G2029R and S217L with Piezo1-GFP had a large effect on the stretch evoked currents, with Y2022A affecting them to a lesser extent (Fig. 8A-C). The stretch evoked activity was seen to be correlated with the extent of fully glycosylated protein, as seen for L939M and G2029R (Fig. 7B). Consistent with all the previous data, G2029R, S217L showed a reduced upper FG band in the Western blot (Fig. 8E). Moreover, co-expression of the G2029R, S217L mutant with the WT protein also reduced the FG band compared with the control (Fig. 8E-G). This finding follows the maximal amount of current elicited from cell-attached patches and suggested a dominant-negative effect (Fig. 8H).

**Figure 8.**
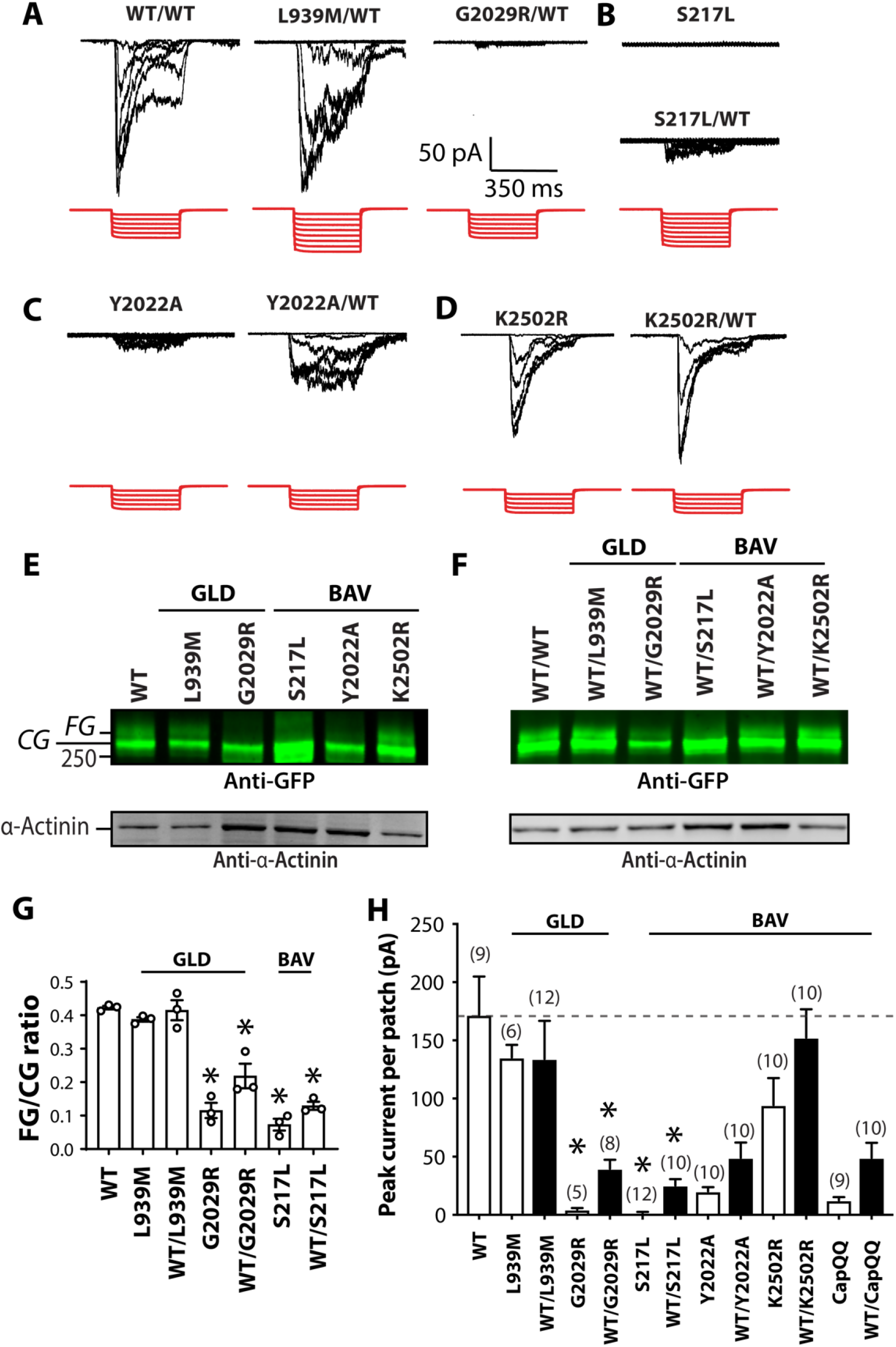
N-linked glycosylation status of Piezo1 in disease-linked variants co-expressed with WT Piezo1. (A) Electrophysiological recordings of Piezo1^-/-^ HEK293T expressing Piezo1-GFP co-expressed with Piezo1-GFP, L939M, G2029R in the cell-attached configuration in response to negative pressure applied using a high-speed pressure-clamp (red). (B) Electrophysiological recordings of bicuspid aortic valve (BAV) linked mutant S217L alone and co-expressed with Piezo1-GFP. (C) Electrophysiological recordings of Y2022A alone and co-expressed with Piezo1-GFP. (D) Electrophysiological recordings of K2502R, alone and co-expressed with Piezo1-GFP. (E) Representative Western blot of Piezo1-GFP, L939M, G2029R, S217L, Y2022A and K2502R. (F) Representative Western blot of Piezo1-GFP/Piezo1-GFP, L939M/Piezo1-GFP, G2029R/Piezo1-GFP, S217L/Piezo1-GFP, Y2022A/Piezo1-GFP and K2502R/Piezo1-GFP. (G) Quantification of upper FG band/lower CG band ratio of Piezo1-GFP, L939M, G2029R, S217L, and L939M, G2029R, S217L co-expressed with Piezo1-GFP. (H) Quantification of peak current elicited per patch of Piezo1-GFP, L939M, G2029R, S217L, Y2022A, K2502R, CapQQ and each mutant co-expressed with Piezo1-GFP. * denotes statistical significance p<0.05 determined by Kruskal-Wallis test with Dunn’s post-hoc test. (GLD – generalized lymphatic dysplasia; BAV – bicuspid aortic valve).

We could not guarantee that 100% of the cells were transfected with *both* WT and mutant protein DNA as they were both GFP fused. However, the patch clamp observations shown in Fig 8 were seen to be consistent by co-expressing a GFP fused Piezo1 with mCherry fused mutant proteins (Fig. 9 & SI Fig. 7). This allowed us to select only cells that were expressing both fused proteins for electrophysiological analysis; almost identical current patterns were recorded (SI Fig. 7) to those of the maximal current data shown in Fig 8H.

**Figure 9.**
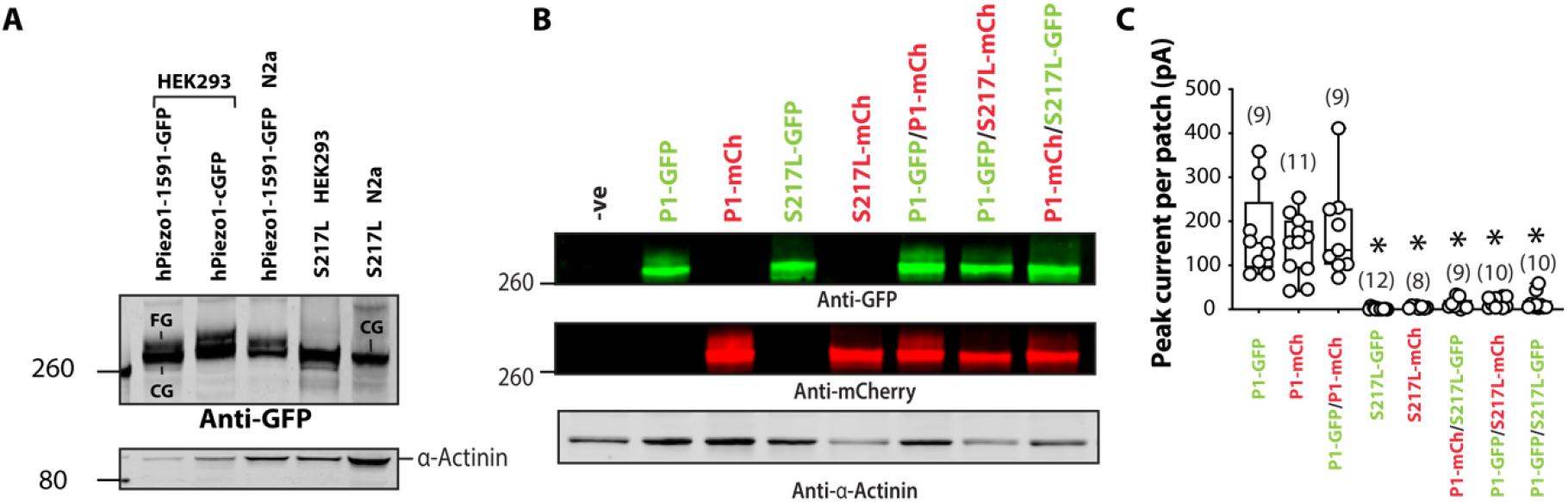
Comprehensive analysis of co-expression of S217L Piezo1 and WT Piezo1 fused to GFP and mCherry. (A) Representative Western blot of Piezo1-GFP and S217L expressed in HEK293 and Neuro2a cells (N2a). (B) Representative Western blot of Piezo1-GFP, Piezo1-mCherry, S217L-GFP, S217L-mCherry, and co-expression of Piezo1-GFP/Piezo1-mCherry, Piezo1-GFP/S217L-mCherry, Piezo1-mCherry/ S217L-GFP. The GFP and mCherry signal are shown separately. (C) Quantification of peak current elicited per patch of all combinations expressed in Piezo1^-/-^ HEK293T cells. * denotes statistical significance p<0.05 determined by Kruskal-Wallis test with Dunn’s post-hoc test.

As final evidence that some mutants displayed a dominant-negative effect we focussed on the S217L mutant. Western blots of this mutant had a single band with little sign of the higher molecular weight fully glycosylated species, regardless of the cell type in which it was expressed (Fig. 8E; Fig. 9A-B). Western blots on cell lysate from Piezo1^-/-^ HEK293T co-transfected with every combination of WT and S217L-GFP and -mCherry fused proteins are shown in Fig 9. Both S217L-GFP and -mCherry fused Piezo1 showed a single band with elimination of the band corresponding to the glycosylated species of the co-transfected wild-type protein (Fig. 9B). The lack of fully glycosylated species correlated well with the stretch evoked current (Fig. 9C).

## Discussion

Post-translational modification is critical for function and localization of transmembrane proteins. N-linked glycosylation is one of the most frequently encountered and heterogeneous forms of co-and post-translational modification. Here we showed that the mechanically-gated ion channel Piezo1 underwent N-linked glycosylation and migrated as a doublet on a Western blot similar to other ion channel proteins like K_v11.1_^18,25^. This double band appearance on Western blots was not dependent on the nature of the molecular tag attached to it. The fully glycosylated species was evident in GFP tagged proteins (C-terminal or 1591 position), mCherry tagged proteins (1591 position), mouse Piezo1 fused to TdTomato and un-tagged natively expressed Piezo1 in fibroblasts.

By using PNGaseF, which specifically cleaves N-linked oligosaccharides, the upper band of human Piezo1 was found to be heavily N-glycosylated (~25 kDa). Treatment with a mixture of deglycosylases produced a similar effect on Western blots implying that the upper band species was unlikely to contain significant amounts of cleavable O-linked oligosaccharides. This was supported by the use of GnT1^-/-^ HEK293S cells. Unlike most cell types this cell has a genetic deletion that prevents the processing of higher order glycans. Western blots from lysates of these cells also lacked an upper band. Using brefeldin A treatment, which inhibits vesicular transport between the ER and Golgi, we showed that the higher order N-glycans were added in the Golgi.

In other ion channel proteins the fully glycosylated protein (upper band) has been used as an indicator of normal membrane trafficking, whereas the core glycosylated protein (lower band) largely represents the immature version present mainly in the ER and Golgi^18,22,53^. Consistent with this knowledge, using unroofed fibroblasts and biotinylation we showed that the N-glycosylated version of natively expressed Piezo1 constituted the major component of the membrane pool of Piezo1. In RBCs only a single Piezo1 band was evident in Western blots and the single band size was reduced on PNGaseF treatment. This finding was consistent with the lack of ER and Golgi in mature RBCs.

The higher order glycosylation of Piezo1 was dependent on two critical Asn sites in the cap region (Fig. 10). Which of these two asparagines that became glycosylated seemed of little consequence as single mutants (N2294Q or N2331Q) could still traffic and produce mechanically evoked currents. A previous mass spectrometry study suggested at least one of these two Asn residues in Piezo1 was glycosylated^43^ which is consistent with a Cryo-EM structure of Piezo2 where one glycosylated asparagine was resolved in the cap^44^. When both residues were ablated (N2294Q and N2331Q) little to no current was present in electrophysiological experiments; and the protein lacked the higher molecular weight band indicative of a protein with higher order N-glycosylation. Interestingly, PNGaseF reduced the size of the core glycosylated N2294Q/N2331Q double mutant protein, thus indicating that other sites also undergo N-linked glycosylation.

**Figure 10.**
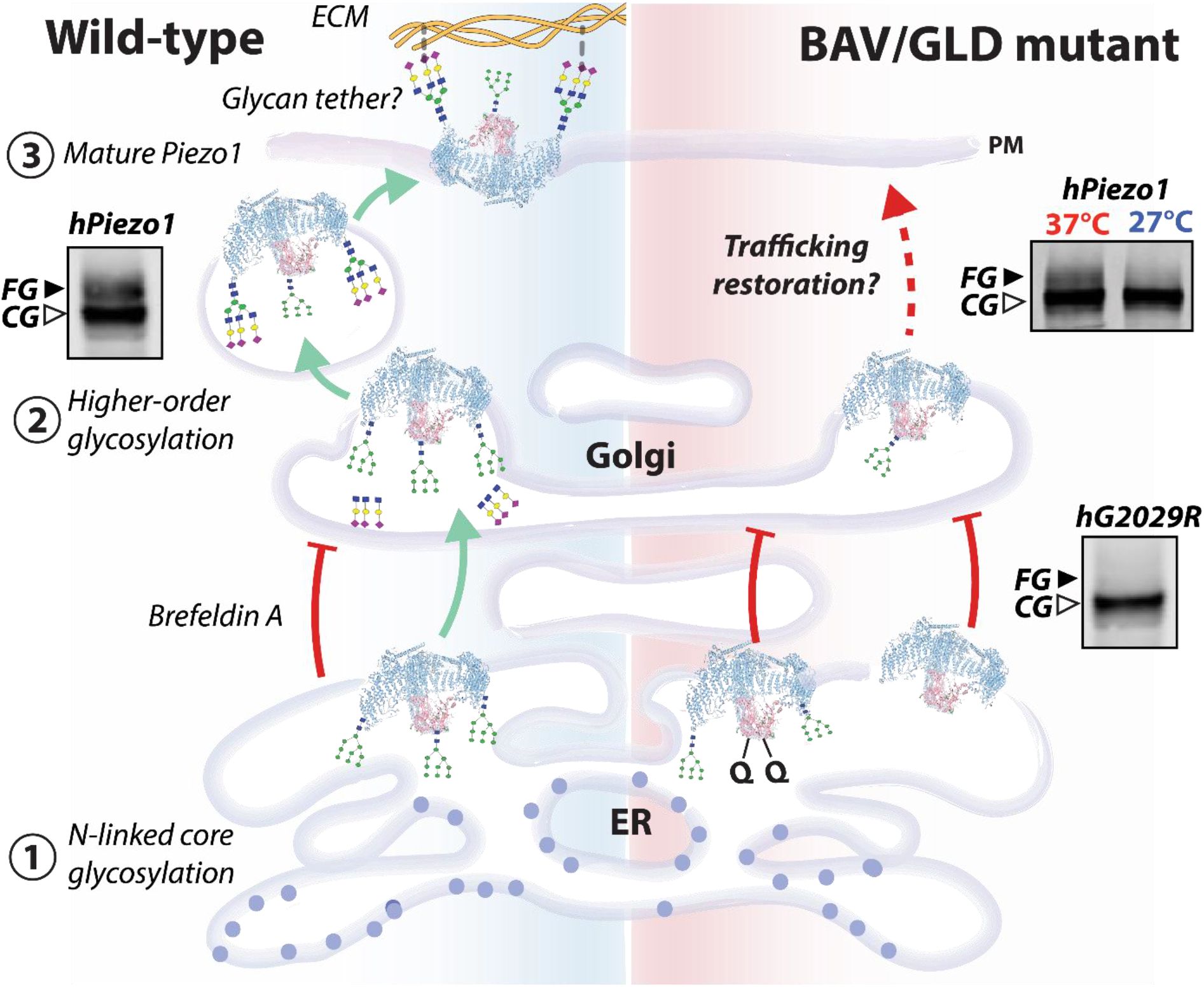
Summary of Piezo1 biosynthetic pathway. Core-glycans added in the cap (N2294/N2331) while folding in the ER are critical for higher order glycans being added in the propeller (primarily at N885) in the Golgi. Aberration of glycosylation impairs trafficking, and perhaps the larger glycans may act as molecular tethers linking to the ECM as suggested for ENaC^62^. Summary adapted from Kanner *et al*.,^66^.

Using a split Piezo1 protein, we showed that both Asn residues became glycosylated and indeed that these residues dictated higher order glycosylation in the propeller regions. Of the six sites predicted to undergo N-glycosylation in the propellers, we identified two crucial residues in the that are the sites of higher order glycosylation (N658 and N885). The analogous residues to N658 and N885 in mouse Piezo1 are present in two loops that were previously identified by the Xiao laboratory to be essential for mouse Piezo1 function; and in electrophysiological stretch assays they gave minimal current consistent with our results^32^.

Thus, our data suggested that the lack of a higher molecular weight species on a Western blot, as seen with the double mutant (N2294Q/N2331Q), indicates aberrant trafficking of Piezo1. This mirrors perfectly what has been reported with K_v11.1_ and CFTR channels^18^. Ultimately it is core N-glycosylation in the cap and propellers that are necessary for trafficking, as current can still be produced in specialized GnT1^-/-^ cells that *cannot* process higher order glycans (although they can produce lower molecular weight mannose containing glycans). In all cells expressing GnT1 that we have tested, such processing in the ER and Golgi results in higher order glycosylation and the upper band on a Western blot.

Probing N-glycosylation status in this manner provides a rapid and reliable method to determine if human Piezo1 variants, that are generated for structure-function studies or in studies of disease-linked variants, exhibit aberrant trafficking exemplified by our data on G2029R, a known trafficking defective Piezo1 variant^28^. We also used glycosylation status to attempt to rescue G2029R using two widely used strategies; low temperature treatment and a pharmacological chaperone^22^. Neither aided G2029R trafficking; but for K_v11.1_ channels only a subset of mutants could be rescued^22^ by the respective treatments. Nevertheless, it is plausible that other Piezo1 mutants may be rescuable using alternative approaches^54^ or pharmacological chaperones.

N-glycosylation status can also be used in co-expression studies to interrogate the impact of disease-linked variants on the WT protein, thus mimicking heterozygosity. Here, we provide evidence that some Piezo1 variants had a dominant negative effect as they reduced WT function. The most notable example of this was S217L. Its effect was very similar to that of dominant negative mutants such as A561V on the trafficking of the K_v11.1_ channel^55^. While there was no guarantee that every cell was expressing exactly the same quantity of mutant and WT Piezo1 proteins, as would be in the case *in vivo*, the data from Western blots and patch clamp analysis were consistent with a dominant negative effect. This was congruent with the fact that the higher molecular weight species (the FG Piezo1 protein) was indicative of the functional membrane protein pool.

While we concluded that the latter mutants are aberrantly trafficked we could not rule out the fact that additional disease-causing mechanisms such as reduced stability or gating phenotypes (particularly for S217L^29^) existed. However our data demonstrated that trafficking deficient Piezo1 mutants were differentially processed in heterologous expression systems as seen in the extensive studies of K_v11.1_^18,49,56–59^ and CFTR^20,60,61^.

We have described this larger species evident on Western blots as the “fully glycosylated” protein which has “higher order” glycosylation. The larger species could have complex glycosylation or hybrid glycans^15^. The exact composition of the glycans is beyond the scope of this study. However, it is likely of interest going forward when trying to decipher whether these glycans may interact with the extracellular matrix and function as molecular tethers as suggested for the epithelial sodium channel (Fig. 10)^62^. The glycans may also facilitate interactions with binding partners such as PECAM1^63^ or E-cadherin^64^, or with specific lipids such as glycolipids^65^.

In integrins, N-glycans are important for clustering. Hence future studies could explore whether higher order Piezo1 glycosylation is involved in the previously identified clustering of Piezo1 in the plasma membrane^9^. If N-glycans are a means of interactions with other membrane proteins, or the ECM^62^, this gives cells an extra dynamic mechanism to regulate mechanosensitivity. This may explain the differential sensitivity to applied force seen when Piezo1 was expressed in HEK293S GnT1^-/-^ cells that cannot process higher order glycans.

In conclusion, we have shown that the fully glycosylated version of Piezo1 *in vitro* is indicative of the mature species, which is localized to the plasma membrane. Thus N-glycosylation status will be valuable in studies of disease-linked variants of Piezo1; and the cell-biological protocols developed here could provide an analytical platform for identifying molecular chaperones to be used in the clinical treatment of Piezo1 trafficking defective variants.

## Acknowledgements

CDC is supported by an NSW Health EMCR Fellowship. CDC and PWK were supported by Australian Research Council Discovery Project Grant DP190100500. The experiments were in part supported by the Victor Chang Cardiac Research Institute Innovation Centre, funded by the NSW Government.

## References

1 Coste, B. et al. Piezo1 and Piezo2 are essential components of distinct mechanically activated cation channels. Science 330, 55–60, doi:10.1126/science.1193270 (2010).

2 Coste, B. et al. Piezo proteins are pore-forming subunits of mechanically activated channels. Nature 483, 176–181, doi:10.1038/nature10812 (2012).

3 Douguet, D., Patel, A., Xu, A., Vanhoutte, P. M. & Honore, E. Piezo Ion Channels in Cardiovascular Mechanobiology. Trends Pharmacol Sci 40, 956–970, doi:10.1016/j.tips.2019.10.002 (2019).

4 Beech, D. J. & Kalli, A. C. Force Sensing by Piezo Channels in Cardiovascular Health and Disease. Arterioscler Thromb Vasc Biol 39, 2228–2239, doi:10.1161/ATVBAHA.119.313348 (2019).

5 Syeda, R. et al. Piezo1 Channels Are Inherently Mechanosensitive. Cell reports 17, 1739–1746, doi:10.1016/j.celrep.2016.10.033 (2016).

6 Cox, C. D. et al. Removal of the mechanoprotective influence of the cytoskeleton reveals PIEZO1 is gated by bilayer tension. Nature communications 7, 10366, doi:10.1038/ncomms10366 (2016).

7 Lewis, A. H. & Grandl, J. Mechanical sensitivity of Piezo1 ion channels can be tuned by cellular membrane tension. eLife 4, doi:10.7554/eLife.12088 (2015).

8 Romero, L. O. et al. Dietary fatty acids fine-tune Piezo1 mechanical response. Nature communications 10, 1200, doi:10.1038/s41467-019-09055-7 (2019).

9 Ridone, P. et al. Disruption of membrane cholesterol organization impairs the activity of PIEZO1 channel clusters. Journal of General Physiology, doi:In press (2020).

10 Tsuchiya, M. et al. Cell surface flip-flop of phosphatidylserine is critical for PIEZO1-mediated myotube formation. Nature communications 9, 2049, doi:10.1038/s41467-018-04436-w (2018).

11 Shi, J. et al. Sphingomyelinase Disables Inactivation in Endogenous PIEZO1 Channels. Cell reports 33, 108225, doi:10.1016/j.celrep.2020.108225 (2020).

12 Cox, C. D. & Gottlieb, P. A. Amphipathic molecules modulate PIEZO1 activity. Biochemical Society transactions 47, 1833–1842, doi:10.1042/BST20190372 (2019).

13 Tannous, A., Pisoni, G. B., Hebert, D. N. & Molinari, M. N-linked sugar-regulated protein folding and quality control in the ER. Seminars in cell & developmental biology 41, 79–89, doi:10.1016/j.semcdb.2014.12.001 (2015).

14 Spiro, R. G. Protein glycosylation: nature, distribution, enzymatic formation, and disease implications of glycopeptide bonds. Glycobiology 12, 43R–56R, doi:10.1093/glycob/12.4.43r (2002).

15 Aebi, M. N-linked protein glycosylation in the ER. Biochimica et biophysica acta 1833, 2430–2437, doi:10.1016/j.bbamcr.2013.04.001 (2013).

16 Glozman, R. et al. N-glycans are direct determinants of CFTR folding and stability in secretory and endocytic membrane traffic. The Journal of cell biology 184, 847–862, doi:10.1083/jcb.200808124 (2009).

17 Gong, Q., Anderson, C. L., January, C. T. & Zhou, Z. Role of glycosylation in cell surface expression and stability of HERG potassium channels. American journal of physiology. Heart and circulatory physiology 283, H77–84, doi:10.1152/ajpheart.00008.2002 (2002).

18 Vandenberg, J. I. et al. hERG K(+) channels: structure, function, and clinical significance. Physiological reviews 92, 1393–1478, doi:10.1152/physrev.00036.2011 (2012).

19 Petrecca, K., Atanasiu, R., Akhavan, A. & Shrier, A. N-linked glycosylation sites determine HERG channel surface membrane expression. The Journal of physiology 515 (Pt 1), 41–48, doi:10.1111/j.1469-7793.1999.041ad.x (1999).

20 Chang, X. B. et al. Role of N-linked oligosaccharides in the biosynthetic processing of the cystic fibrosis membrane conductance regulator. Journal of cell science 121, 2814–2823, doi:10.1242/jcs.028951 (2008).

21 Cai, Y. et al. Altered trafficking and stability of polycystins underlie polycystic kidney disease. The Journal of clinical investigation 124, 5129–5144, doi:10.1172/JCI67273 (2014).

22 Anderson, C. L. et al. Large-scale mutational analysis of Kv11.1 reveals molecular insights into type 2 long QT syndrome. Nature communications 5, 5535, doi:10.1038/ncomms6535 (2014).

23 Ke, Y. et al. Trafficking defects in PAS domain mutant Kv11.1 channels: roles of reduced domain stability and altered domain-domain interactions. The Biochemical journal 454, 69–77, doi:10.1042/BJ20130328 (2013).

24 Foo, B., Williamson, B., Young, J. C., Lukacs, G. & Shrier, A. hERG quality control and the long QT syndrome. The Journal of physiology 594, 2469–2481, doi:10.1113/JP270531 (2016).

25 Apaja, P. M. et al. Ubiquitination-dependent quality control of hERG K+ channel with acquired and inherited conformational defect at the plasma membrane. Molecular biology of the cell 24, 3787–3804, doi:10.1091/mbc.E13-07-0417 (2013).

26 Perry, M. D. et al. Rescue of protein expression defects may not be enough to abolish the pro-arrhythmic phenotype of long QT type 2 mutations. The Journal of physiology 594, 4031–4049, doi:10.1113/JP271805 (2016).

27 Fotiou, E. et al. Novel mutations in PIEZO1 cause an autosomal recessive generalized lymphatic dysplasia with non-immune hydrops fetalis. Nature communications 6, 8085, doi:10.1038/ncomms9085 (2015).

28 Lukacs, V. et al. Impaired PIEZO1 function in patients with a novel autosomal recessive congenital lymphatic dysplasia. Nature communications 6, 8329, doi:10.1038/ncomms9329 (2015).

29 Faucherre, A. et al. Piezo1 is required for outflow tract and aortic valve development. Journal of molecular and cellular cardiology 143, 51–62, doi:10.1016/j.yjmcc.2020.03.013 (2020).

30 Saotome, K. et al. Structure of the mechanically activated ion channel Piezo1. Nature 554, 481–486, doi:10.1038/nature25453 (2018).

31 Guo, Y. R. & MacKinnon, R. Structure-based membrane dome mechanism for Piezo mechanosensitivity. eLife 6, doi:10.7554/eLife.33660 (2017).

32 Zhao, Q. et al. Structure and mechanogating mechanism of the Piezo1 channel. Nature 554, 487–492, doi:10.1038/nature25743 (2018).

33 Bae, C., Gnanasambandam, R., Nicolai, C., Sachs, F. & Gottlieb, P. A. Xerocytosis is caused by mutations that alter the kinetics of the mechanosensitive channel PIEZO1. Proceedings of the National Academy of Sciences of the United States of America 110, E1162–1168, doi:10.1073/pnas.1219777110 (2013).

34 Biazik, J., Yla-Anttila, P., Vihinen, H., Jokitalo, E. & Eskelinen, E. L. Ultrastructural relationship of the phagophore with surrounding organelles. Autophagy 11, 439–451, doi:10.1080/15548627.2015.1017178 (2015).

35 Kuo, J. C., Han, X., Yates, J. R., 3rd & Waterman, C. M. Isolation of focal adhesion proteins for biochemical and proteomic analysis. Methods in molecular biology 757, 297–323, doi:10.1007/978-1-61779-166-6_19 (2012).

36 Ranade, S. S. et al. Piezo1, a mechanically activated ion channel, is required for vascular development in mice. Proceedings of the National Academy of Sciences of the United States of America 111, 10347–10352, doi:10.1073/pnas.1409233111 (2014).

37 Nebenfuhr, A., Ritzenthaler, C. & Robinson, D. G. Brefeldin A: deciphering an enigmatic inhibitor of secretion. Plant physiology 130, 1102–1108, doi:10.1104/pp.011569 (2002).

38 Gordon, S. E., Munari, M. & Zagotta, W. N. Visualizing conformational dynamics of proteins in solution and at the cell membrane. eLife 7, doi:10.7554/eLife.37248 (2018).

39 Taraska, J. W. A primer on resolving the nanoscale structure of the plasma membrane with light and electron microscopy. The Journal of general physiology 151, 974–985, doi:10.1085/jgp.201812227 (2019).

40 Yao, M. et al. Force-dependent Piezo1 recruitment to focal adhesions regulates adhesion maturation and turnover specifically in non-transformed cells (Cold Spring Harbor Laboratory, 2020).

41 Ellefsen, K. L. et al. Myosin-II mediated traction forces evoke localized Piezo1-dependent Ca(2+) flickers. Commun Biol 2, 298, doi:10.1038/s42003-019-0514-3 (2019).

42 Blom, N., Sicheritz-Ponten, T., Gupta, R., Gammeltoft, S. & Brunak, S. Prediction of post-translational glycosylation and phosphorylation of proteins from the amino acid sequence. Proteomics 4, 1633–1649, doi:10.1002/pmic.200300771 (2004).

43 Wollscheid, B. et al. Mass-spectrometric identification and relative quantification of N-linked cell surface glycoproteins. Nature biotechnology 27, 378–386, doi:10.1038/nbt.1532 (2009).

44 Wang, L. et al. Structure and mechanogating of the mammalian tactile channel PIEZO2. Nature 573, 225–229, doi:10.1038/s41586-019-1505-8 (2019).

45 Lewis, A. H. & Grandl, J. Inactivation Kinetics and Mechanical Gating of Piezo1 Ion Channels Depend on Subdomains within the Cap. Cell reports 30, 870–880 e872, doi:10.1016/j.celrep.2019.12.040 (2020).

46 Struwe, W. B. & Robinson, C. V. Relating glycoprotein structural heterogeneity to function - insights from native mass spectrometry. Current opinion in structural biology 58, 241–248, doi:10.1016/j.sbi.2019.05.019 (2019).

47 Botello-Smith, W. M. et al. A mechanism for the activation of the mechanosensitive Piezo1 channel by the small molecule Yoda1. Nature communications 10, 4503, doi:10.1038/s41467-019-12501-1 (2019).

48 Bae, C., Suchyna, T. M., Ziegler, L., Sachs, F. & Gottlieb, P. A. Human PIEZO1 Ion Channel Functions as a Split Protein. PloS one 11, e0151289, doi:10.1371/journal.pone.0151289 (2016).

49 Wang, Y. et al. The role and mechanism of chaperones Calnexin/Calreticulin in which ALLN selectively rescues the trafficking defective of HERG-A561V mutation. Biosci Rep 38, doi:10.1042/BSR20171269 (2018).

50 Wang, X., Koulov, A. V., Kellner, W. A., Riordan, J. R. & Balch, W. E. Chemical and biological folding contribute to temperature-sensitive DeltaF508 CFTR trafficking. Traffic (Copenhagen, Denmark) 9, 1878–1893, doi:10.1111/j.1600-0854.2008.00806.x (2008).

51 Zheng, W., Nikolaev, Y. A., Gracheva, E. O. & Bagriantsev, S. N. Piezo2 integrates mechanical and thermal cues in vertebrate mechanoreceptors. Proceedings of the National Academy of Sciences, 201910213, doi:10.1073/pnas.1910213116 (2019).

52 Evans, E. L. et al. Yoda1 analogue (Dooku1) which antagonizes Yoda1-evoked activation of Piezo1 and aortic relaxation. British journal of pharmacology 175, 1744–1759, doi:10.1111/bph.14188 (2018).

53 Ke, Y., Hunter, M. J., Ng, C. A., Perry, M. D. & Vandenberg, J. I. Role of the cytoplasmic N-terminal Cap and Per-Arnt-Sim (PAS) domain in trafficking and stabilization of Kv11.1 channels. The Journal of biological chemistry 289, 13782–13791, doi:10.1074/jbc.M113.531277 (2014).

54 Kanner, S. A., Shuja, Z., Choudhury, P., Jain, A. & Colecraft, H. M. Targeted deubiquitination rescues distinct trafficking-deficient ion channelopathies. Nature methods, doi:10.1038/s41592-020-00992-6 (2020).

55 Kagan, A., Yu, Z., Fishman, G. I. & McDonald, T. V. The dominant negative LQT2 mutation A561V reduces wild-type HERG expression. The Journal of biological chemistry 275, 11241–11248, doi:10.1074/jbc.275.15.11241 (2000).

56 Foo, B. et al. Mutation-specific peripheral and ER quality control of hERG channel cell surface expression. Scientific reports 9, 6066, doi:10.1038/s41598-019-42331-6 (2019).

57 Smith, J. L. et al. Pharmacological correction of long QT-linked mutations in KCNH2 (hERG) increases the trafficking of Kv11.1 channels stored in the transitional endoplasmic reticulum. American journal of physiology. Cell physiology 305, C919–930, doi:10.1152/ajpcell.00406.2012 (2013).

58 Smith, J. L. et al. Trafficking-deficient hERG K(+) channels linked to long QT syndrome are regulated by a microtubule-dependent quality control compartment in the ER. American journal of physiology. Cell physiology 301, C75–85, doi:10.1152/ajpcell.00494.2010 (2011).

59 Zhou, Z., Gong, Q. & January, C. T. Correction of defective protein trafficking of a mutant HERG potassium channel in human long QT syndrome. Pharmacological and temperature effects. The Journal of biological chemistry 274, 31123–31126, doi:10.1074/jbc.274.44.31123 (1999).

60 Loo, M. A. et al. Perturbation of Hsp90 interaction with nascent CFTR prevents its maturation and accelerates its degradation by the proteasome. The EMBO journal 17, 6879–6887, doi:10.1093/emboj/17.23.6879 (1998).

61 Owsianik, G., Cao, L. & Nilius, B. Rescue of functional DeltaF508-CFTR channels by co-expression with truncated CFTR constructs in COS-1 cells. FEBS letters 554, 173–178 (2003).

62 Knoepp, F. et al. Shear force sensing of epithelial Na(+) channel (ENaC) relies on N-glycosylated asparagines in the palm and knuckle domains of alphaENaC. Proceedings of the National Academy of Sciences of the United States of America 117, 717–726, doi:10.1073/pnas.1911243117 (2020).

63 Chuntharpursat-Bon, E. et al. Cell adhesion molecule interaction with Piezo1 channels is a mechanism for sub cellular regulation of mechanical sensitivity. 602532, doi:10.1101/602532 %J bioRxiv (2019).

64 Wang, J., Jiang, J., Yang, X., Wang, L. & Xiao, B. Tethering Piezo channels to the actin cytoskeleton for mechanogating via the E-cadherin-β-catenin mechanotransduction complex. 2020.2005.2012.092148, doi:10.1101/2020.05.12.092148 %J bioRxiv (2020).

65 Buyan, A. et al. Piezo1 Forms Specific, Functionally Important Interactions with Phosphoinositides and Cholesterol. Biophysical journal 119, 1683–1697, doi:10.1016/j.bpj.2020.07.043 (2020).

66 Kanner, S. A., Jain, A. & Colecraft, H. M. Development of a High-Throughput Flow Cytometry Assay to Monitor Defective Trafficking and Rescue of Long QT2 Mutant hERG Channels. Frontiers in physiology 9, 397, doi:10.3389/fphys.2018.00397 (2018).

