## Supplementary Information for "Modified N-linked glycosylation status predicts trafficking defective human Piezo1 channel mutations"

[1]Molecular Cardiology and Biophysics Division, Victor Chang Cardiac Research Institute, Sydney, Australia. [2] St Vincent's Clinical School, Faculty of Medicine, University of New South Wales, Sydney, Australia. [3]Mechanobiology Institute, National University of Singapore, Singapore. [4]School of Biomedical Engineering, Faculty of Engineering, The University of Sydney, Camperdown, New South Wales, Australia. [7]School of Life and Environmental Sciences, University of Sydney, Sydney, New South Wales, Australia.

***Running title:*** N-linked glycosylation in Piezo1 channels

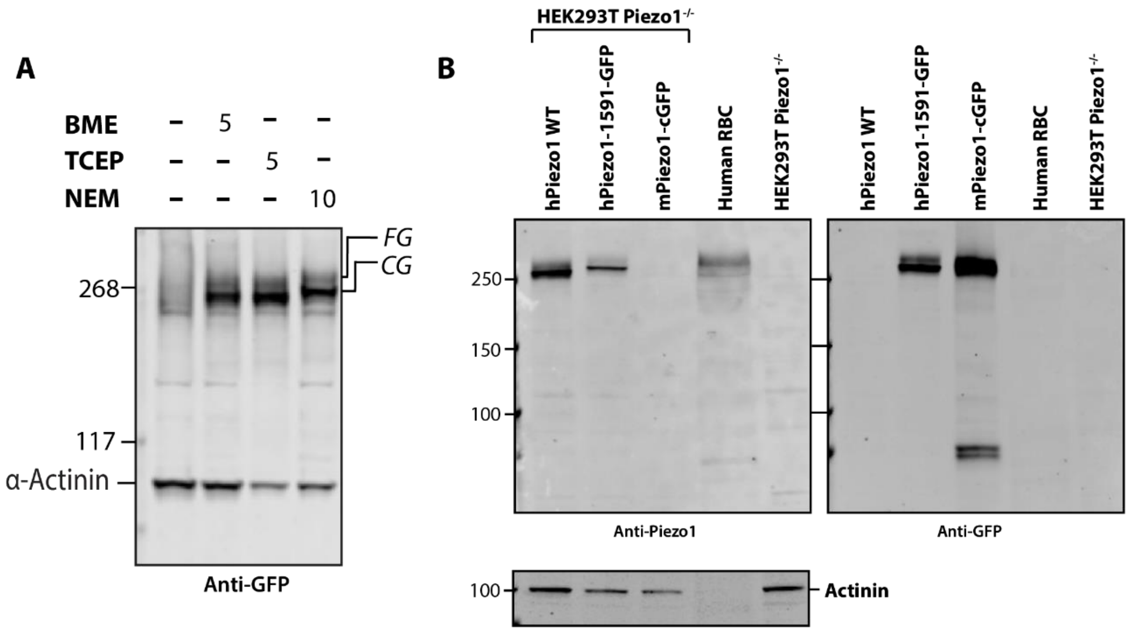


**SI Figure1. Primary monoclonal Piezo1 antibody (Novus Biologicals) reproducibly recognizes human Piezo1 but fails to recognize mouse Piezo1.** (A) ) A representative western blot of Piezo1-1591-GFP fusion lysate with no reducing agent present in the lysis buffer compared with the addition of 5 mM BME, 5 mM TCEP and 10 mM N-ethylmaleimide expressed in Piezo1$^{-/-}$ HEK293T. Notice the doublet on the Western blot in the presence of a reducing agent or alkylating agent. (B) Left panel -Representative Western blot using Novus primary monoclonal (Cat# NBP2-75617, Novus Biologicals) anti-Piezo1 antibody with samples of untagged human Piezo1, Piezo1-1591-GFP fusion and mouse Piezo1-C-terminal-GFP expressed in HEK293T Piezo1$^{-/-}$ compared to human red blood cell lysate and HEK293T Piezo1$^{-/-}$ lysate as a negative control. Right panel- Same blot as that shown in left panel probed with an anti-GFP antibody.

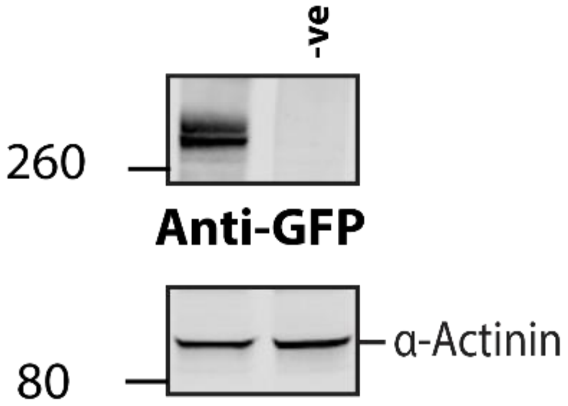


**SI Figure2. Piezo1 expression in Neuro2a also runs as two bands on Western blots.** Representative Western blot of transiently transfected Neuro2a cells with Piezo1-GFP and a vector transfected negative control.

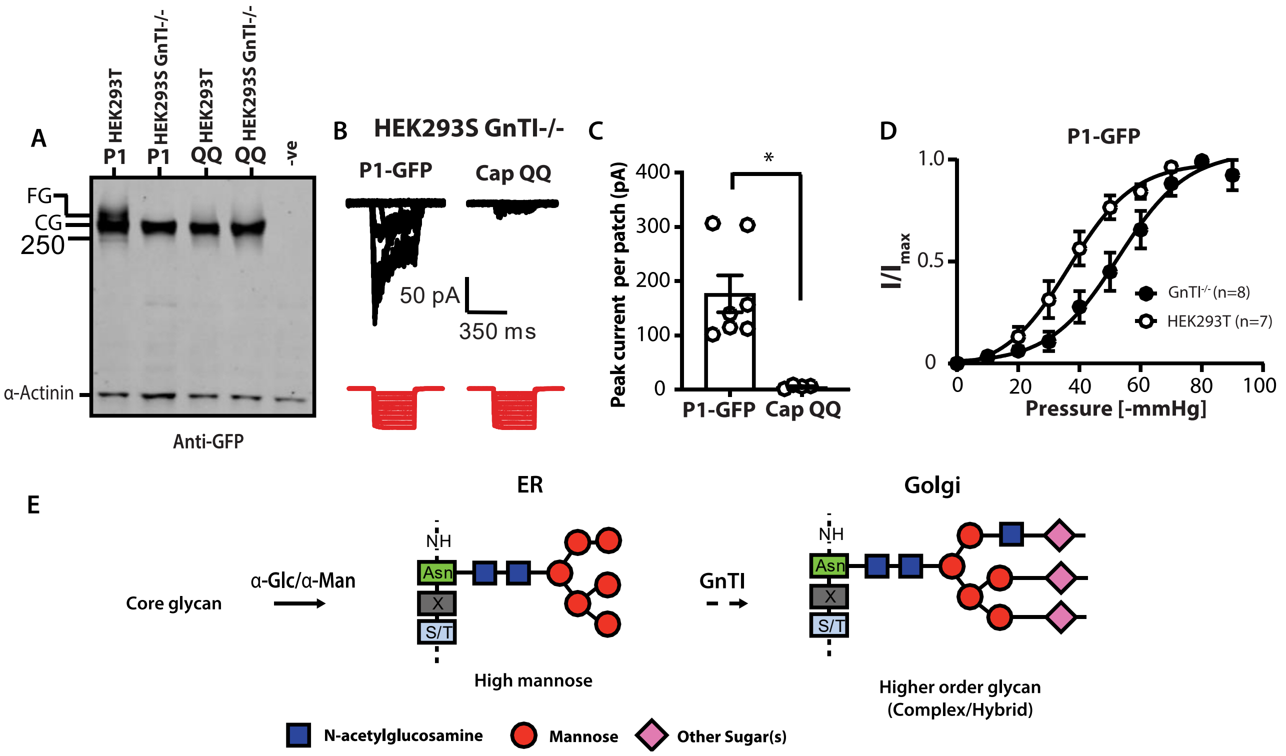


**SI Figure3. Higher order glycosylation does not happen in GnT1-/- HEK293S cells.** (A) Representative Western blot comparing Piezo1-GFP (P1-GFP), CapQQ mutant expressed in HEK293T Piezo1$^{-/-}$ and HEK293S GnT1$^{-/-}$. (B) Electrophysiological recordings of HEK293S GnT1$^{-/-}$ expressing Piezo1-GFP and CapQQ in the cell-attached configuration in response to negative pressure applied using a high-speed pressure-clamp (red). (C) Quantification of peak current elicited per patch of Piezo1-GFP and CapQQ expressed in HEK293S GnT1$^{-/-}$. (D) Pressure response curve of Piezo1-GFP expressed in Piezo1$^{-/-}$ HEK293T compared to Gnt1-/- HEK293S. (E) A cartoon representing N-acetylglucosaminyl-transferase I (GnT1) enzyme processing of N-glycans from high-mannose to higher molecular weight glycans.

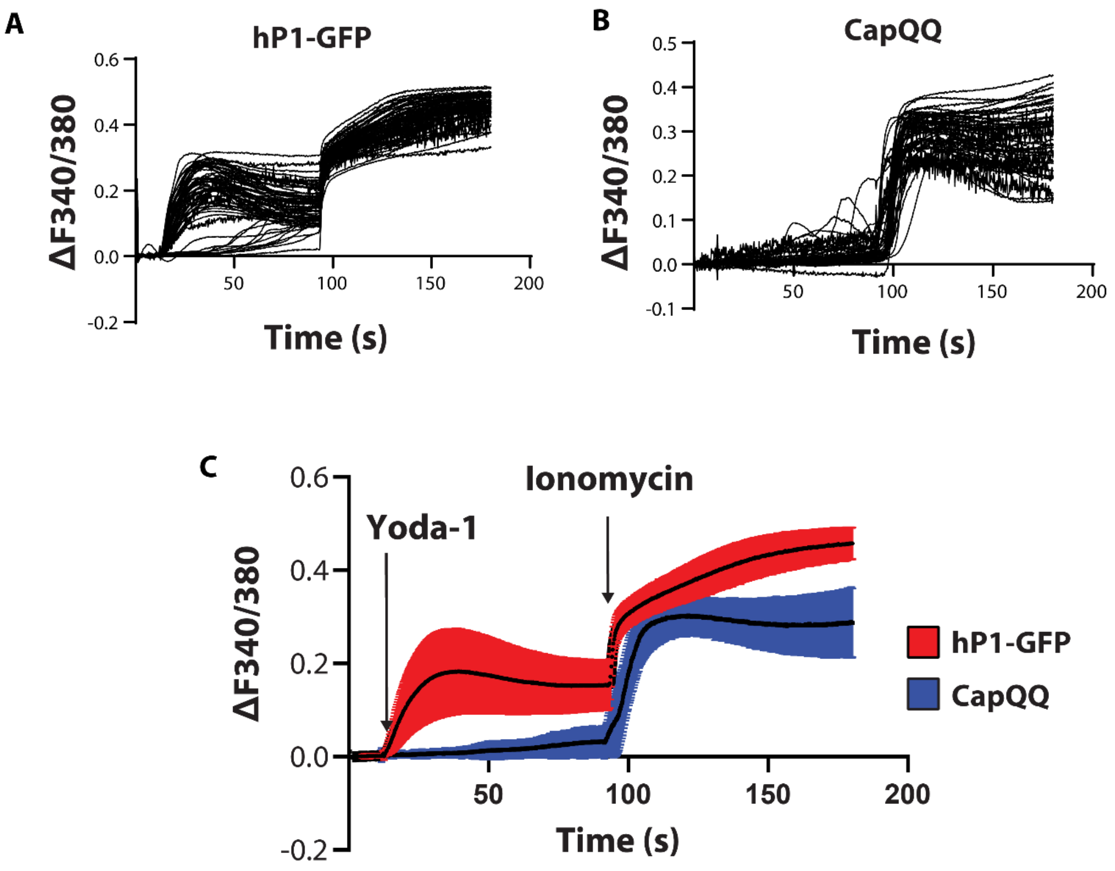


**SI Figure4. Yoda-1 fails to elicit a $Ca^{2+}$ response in HEK293T Piezo1$^{-/-}$ cells transfected with the CapQQ mutant.** (A) Individual cell responses to 2 µM Yoda-1 and 5uM ionomycin for HEK293T Piezo1$^{-/-}$ cells transfected with human Piezo1 (hP1-GFP). (B) Individual cell responses to 2 µM Yoda-1 and 5 µM ionomycin for HEK293T Piezo1$^{-/-}$ cells transfected with CapQQ. (C) Comparison between the response of Piezo1 and the CapQQ mutant to Yoda-1 stimulation (2 µM). Data shows n=70 cells from three independent experiments - mean ± SD.

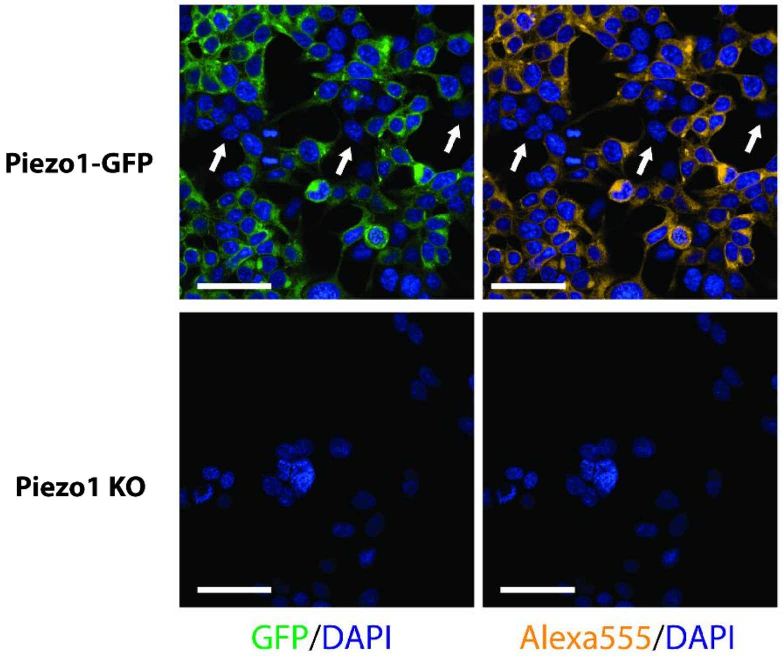


**SI Figure 5. Fidelity of mouse monoclonal anti-Piezo1 antibody for immunofluorescence.** Immunofluorescence of Piezo1$^{-/-}$ HEK293T cells expressing Piezo1-GFP probed using Novus monoclonal anti-Piezo1 primary antibody and its secondary antibody conjugated with Alexa555 fluorophore. Piezo1$^{-/-}$ HEK293T cells are shown as negative control. DAPI signal represents nuclei. Arrows point out un-transfected cells, which has neither GFP nor Alexa555 signal.

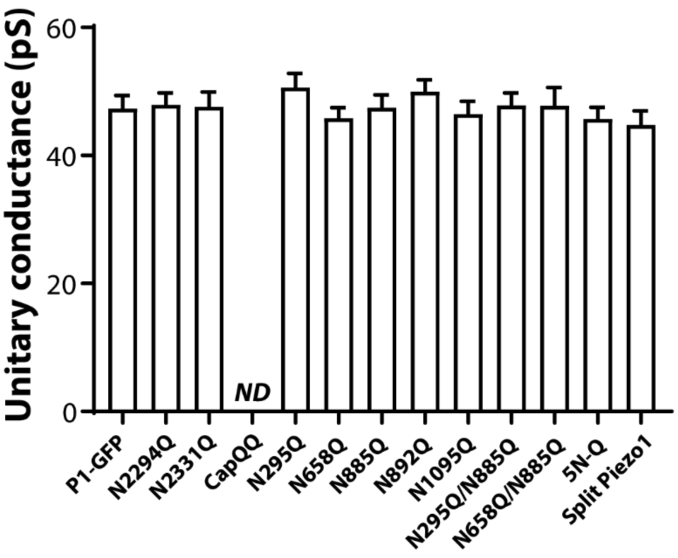


**SI Fig 6. Comparison of unitary conductance of glycosylation mutants.** Unitary conductance of Piezo1 variants indicated in the presence of high extracellular $K^+$ to zero membrane potential. Data represents mean ± SEM; n=5-8.

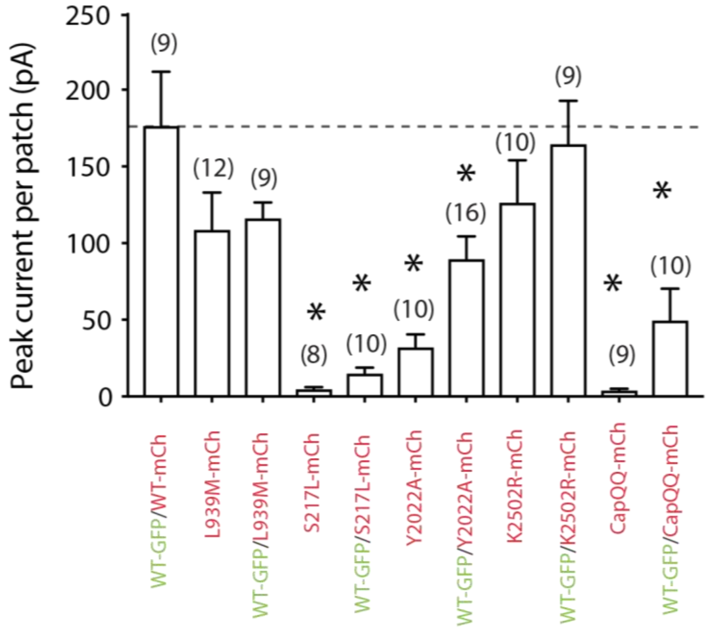


**SI Figure 7. Large scale analysis of co-transfection of Piezo1-GFP with mCherry fused mutants.** Quantification of peak current elicited per patch of Piezo1-GFP/Piezo1-mCherry, L939M-mCherry alone, Piezo1-GFP/L939M-mCherry, S217L- mCherry alone, Piezo1-GFP/S217L-mCherry, Y2022A-mCherry alone, Piezo1-GFP/Y2022A-mCherry, K2502R-mCherry alone, Piezo1-GFP/ K2502R-mCherry, CapQQ-mCherry alone and Piezo1-GFP/CapQQ-mCherry expressed in HEK293T Piezo1$^{-/-}$ cells. * Data represents mean± SEM; P<0.05 determined by Kruskal-Wallis test with Dunn's post-hoc test.
